# The transcriptional corepressor CTBP-1 acts with the SOX family transcription factor EGL-13 to maintain AIA interneuron cell identity in *C. elegans*

**DOI:** 10.1101/2021.07.28.454018

**Authors:** Josh Saul, Takashi Hirose, H. Robert Horvitz

**Author notes:** Sysmex Corporation, 4-4-4 Takatsukadai, Nishi-ku, Kobe 651-2271, Japan (Takashi Hirose). Corresponding author (HRH).

## Abstract

Cell identity is characterized by a distinct combination of gene expression, cell morphology and cellular function established as progenitor cells divide and differentiate. Following establishment, cell identities can be unstable and require active and continuous maintenance throughout the remaining life of a cell. Mechanisms underlying the maintenance of cell identities are incompletely understood. Here we show that the gene *ctbp-1,* which encodes the transcriptional corepressor <u>C</u>-terminal <u>b</u>inding <u>p</u>rotein-1 (CTBP-1), is essential for the maintenance of the identities of the two AIA interneurons in the nematode *Caenorhabditis elegans*. *ctbp-1* is not required for the establishment of the AIA cell fate but rather functions cell-autonomously and can act in later larval stage and adult worms to maintain proper AIA gene expression, morphology and function. From a screen for suppressors of the *ctbp-1* mutant phenotype, we identified the gene *egl-13,* which encodes a SOX family transcription factor. We found that *egl-13* regulates AIA function and aspects of AIA gene expression, but not AIA morphology. We conclude that the CTBP-1 protein maintains AIA cell identity in part by utilizing EGL-13 to repress transcriptional activity in the AIAs. More generally, we propose that transcriptional corepressors like CTBP-1 might be critical factors in the maintenance of cell identities, harnessing the DNA-binding specificity of transcription factors like EGL-13 to selectively regulate gene expression in a cell-specific manner.

## Introduction

Over the course of animal development, complex networks of transcription factors act and interact to drive the division and differentiation of progenitor cells towards terminal cell identities [1–8]. These networks of transcriptional activity often culminate in the activation of master transcriptional regulators that are responsible for directing the differentiation of a diverse range of cell and tissue types [4,9–12]. Examples of such master transcriptional regulators include the mammalian bHLH transcription factor MyoD, which specifies skeletal muscle cells [13–15]; the *Drosophila* Pax-family transcription factor Eyeless, which drives differentiation of the fly eye [16–20]; and the *C. elegans* GATA transcription factor ELT-2, essential for development of the worm intestine [21–24]. Many such master transcriptional regulators are not only required to establish the identities of specific cell types but are subsequently continuously required to maintain those identities for the remaining life of the cell [4,23,25–29]. Defects in the maintenance of cell identities can manifest as late-onset misregulated gene expression, altered morphology or disrupted cellular function, and often become progressively worse as the cell ages [25,28,30–32].

Previous studies of the nematode *Caenorhabditis elegans* have identified a class of master transcriptional regulators, termed terminal selectors [4,12,33– 37]. Terminal selectors drive the expression of whole batteries of gene activity that ultimately define the unique features of many different cell types [10–12]. Individual terminal selectors have been shown to contribute to the establishment and maintenance of multiple distinct *C. elegans* cell types and to drive the expression of many cell-type specific genes [7,37–41]. However, it has been unclear how individual terminal selectors can drive the expression of cell-type specific genes in only the appropriate cell types rather than in all cells in which they act [42–44]. Recent work has shown that terminal selectors appear to broadly activate the expression of many genes, including cell-type specific genes, in all cells in which they function [42, 45]. Piecemeal assemblies of transcription factors are then responsible for pruning this broad expression to restrict expression of cell-type specific genes to the appropriate cell types [42, 45]. This restriction of the activation of gene expression by terminal selectors appears to be an essential aspect of proper cell-identity maintenance [28,42,45,46]. However, it is not known how the myriad of transcription factors utilized to restrict terminal selector gene activation are coordinated and controlled.

Here we report the discovery that the *C. elegans* gene *ctbp-1,* which encodes the sole worm ortholog of the <u>C</u>-terminal <u>B</u>inding <u>P</u>rotein (CtBP) family of transcriptional corepressors [47–55], functions to maintain the cell identity of the two AIA interneurons. We demonstrate that CTBP-1 functions with the SOX-family transcription factor EGL-13 [56, 57] to maintain multiple aspects of the AIA cell identity and propose that CTBP-1 does so in part by utilizing EGL-13 to repress transcriptional activity in the AIAs.

## Results

### Mutations in *ctbp-1* cause *ceh-28* reporter misexpression in the AIA neurons

In previous studies, we screened for and characterized mutations that prevent the programmed cell death of the sister cell of the *C. elegans* M4 neuron [58, 59]. For these screens, we used the normally M4-specific GFP transcriptional reporter *P_ceh-28_::gfp* and identified isolates with an undead M4 sister cell, which expresses characteristics normally expressed by the M4 cell, on the basis of ectopic GFP expression. In addition to mutants with an undead M4 sister cell, we isolated 18 mutant strains that express *P_ceh-28_::gfp* in a manner uncharacteristic of M4 or its undead sister. These mutants express *P_ceh-28_::gfp* in a bilaterally symmetric pair of cells located near the posterior of the *C. elegans* head, far from both M4 and the single M4 sister cell (Fig. 1A).

**Figure 1.**
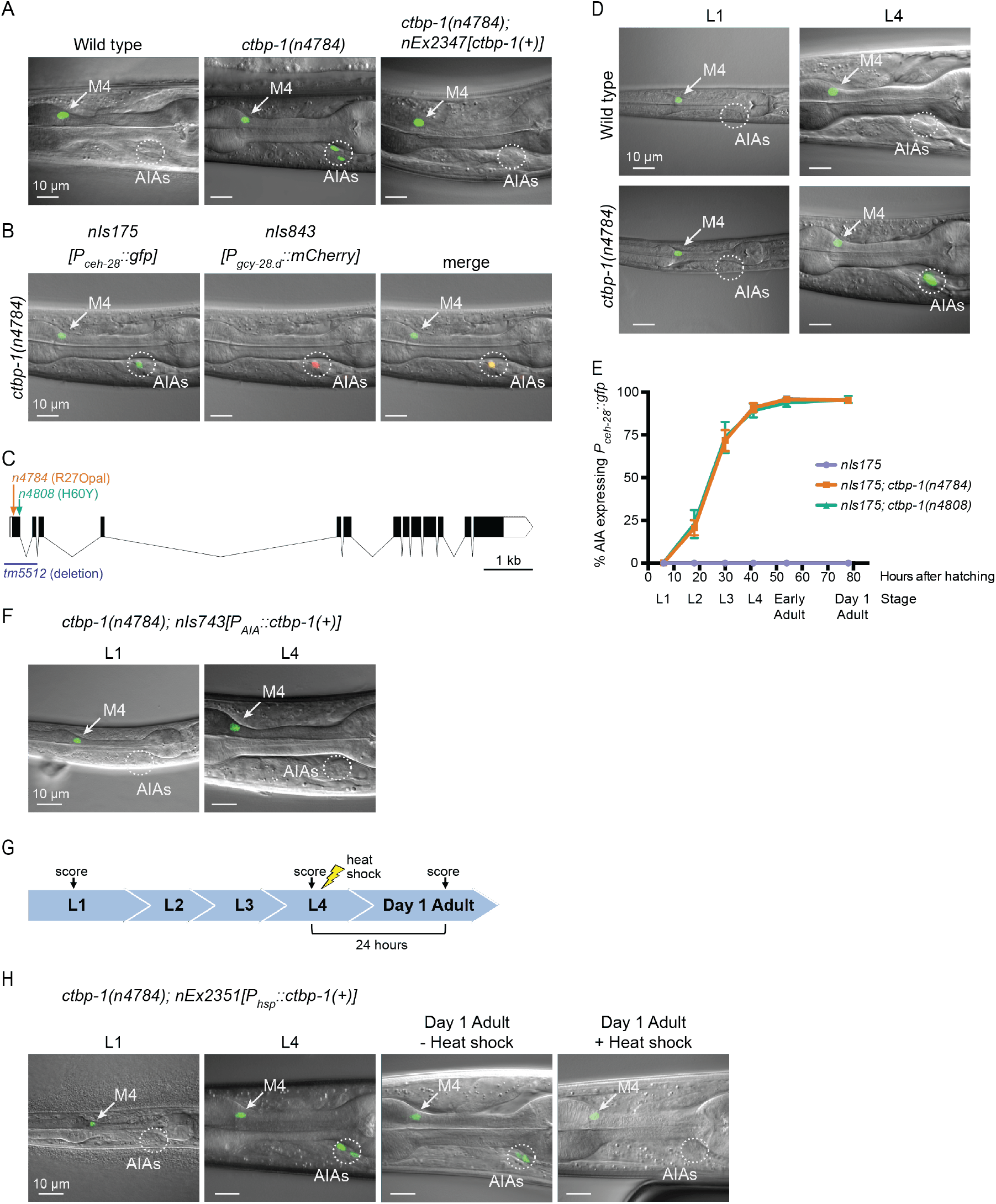
*ctbp-1* mutants misexpress *P_ceh-28_::gfp* in the AIA neurons. (A) Expression of the M4-specific marker *nIs175[P_ceh-28_::gfp]* in the wild type (left panel), a *ctbp-1(n4784)* mutant (middle panel), and a *ctbp-1* mutant carrying an extrachromosomal array expressing wild-type *ctbp-1* under its native promoter (*nEx2347*) (right panel). Arrow, M4 neuron. Circle, AIAs. Scale bar, 10 µm. (B) A *ctbp-1(n4784)* mutant expressing *nIs175* (left panel) and the AIA marker *nIs843[P_gcy-28.d_::mCherry]* (middle panel). Merge, right panel. Arrow, M4 neuron. Circle, AIAs. Scale bar, 10 µm. (C) Gene diagram of the *ctbp-1a* isoform. Arrows (above), point mutations. Line (below), deletion. Scale bar (bottom right), 1 kb. Additional *ctbp-1* alleles are shown in Fig. S1B. (D) *nIs175* expression in wild-type (top) and *ctbp-1(n4784)* (bottom) worms at the L1 larval stage (left) and L4 larval stage (right). Arrow, M4 neuron. Circle, AIAs. Scale bar, 10 µm. (E) Percentage of wild-type, *ctbp-1(n4784)*, and *ctbp-1(n4808)* worms expressing *nIs175* in the AIA neurons over time. Time points correspond to the L1, L2, L3, and L4 larval stages, early adult, and day 1 adult worms (indicated below X axis). Mean ± SEM. *n* ≥ 60 worms scored per strain per stage, 4 biological replicates. (F) Expression of *nIs175* in *ctbp-1* mutants containing a transgene driving expression of wild-type *ctbp-1* under an AIA-specific promoter (*nIs743[P_gcy-28.d_::ctbp-1(+)]*) in L1 and L4 larval worms. Arrow, M4 neuron. Circle, AIAs. Scale bar, 10 µm. (G) Schematic for the heat shock experiment shown in Fig. 1H. (H) *nIs175* expression in *ctbp-1(n4784)* mutants carrying the heat shock-inducible transgene *nEx2351[P_hsp-16.2_::ctbp-1(+);P_hsp-16.41_::ctbp-1(+)]*. Arrow, M4 neuron. Circle, AIAs. Scale bar, 10 µm. All strains shown contain the transgene *nIs175[P_ceh-28_::gfp]*. Images are oriented such that left corresponds to anterior, top to dorsal.

These mutations define a single complementation group and all 18 mutant strains have mutations in the transcriptional corepressor gene *ctbp-1* (Figs. 1C; S1A-B; S2A). These *ctbp-1* alleles include three splice-site mutations and nine nonsense mutations (such as the mutation *n4784,* an early nonsense mutation and one of many presumptive null alleles of the gene). The mutant phenotype is recessive, and a transgenic construct carrying a wild-type copy of *ctbp-1* expressed under its native promoter fully rescued the GFP misexpression caused by *n4784* (Figs. 1A; S2A). *tm5512*, a 632 bp deletion spanning the transcription start site and first two exons of the *ctbp-1a* isoform and a presumptive null allele of this gene [60], likewise caused *P_ceh-28_::gfp* misexpression in two cells in the posterior region of the head (Fig. S1C-D), similar to our *ctbp-1* isolates. These findings demonstrate that loss of *ctbp-1* function is responsible for *P_ceh-28_::gfp* misexpression.

To determine the identity of the cells misexpressing the normally M4-specific marker *P_ceh-28_::gfp*, we examined reporters for cells in the vicinity of the observed misexpression in *ctbp-1* mutants. The AIA-neuron reporter *nIs843[P_gcy-28.d_::mCherry]* showed complete overlap with misexpressed *P_ceh-28_::gfp*, indicating that the cells misexpressing the M4 reporter are the two bilaterally symmetric and embryonically-generated AIA interneurons (Fig. 1B).

### The penetrance of *ceh-28* reporter misexpression in the AIA neurons increases with age

While characterizing *ctbp-1* mutants, we noticed that fewer young worms misexpress *P_ceh-28_::gfp* in the AIAs than do older worms (Fig. 1D). To investigate the temporal aspect of this phenotype, we scored *ctbp-1* mutants for *P_ceh-28_::gfp* misexpression throughout the four worm larval stages (L1-L4) and into the first day of adulthood (“early” and “day 1” adults). *ctbp-1* mutants rarely misexpressed *P_ceh-28_::gfp* at early larval stages, but displayed an increasing penetrance, though invariant expressivity, of this defect as worms transitioned through larval development, such that by the last larval stage (L4) nearly all worms exhibited reporter misexpression specifically and solely in the AIAs (Fig. 1E). A similar stage-dependent increase in reporter expression in *ctbp-1* mutants occurred in mutants carrying a second independently-generated *ceh-28* reporter, *nIs348[P_ceh-28_::mCherry]* (Fig. S1E). These results demonstrate that *ctbp-1* function prevents an age-dependent misexpression of the M4-specific gene *ceh-28* in the unrelated AIA neurons.

We next asked in what cells and at what stages *ctbp-1* functions to suppress *P_ceh-28_::gfp* expression in the AIAs. We generated a transgenic construct that expresses wild-type *ctbp-1* specifically in the AIAs, *nIs743[P_gcy-28.d_::ctbp-1(+)]* (hereafter referred to as *nIs743[P_AIA_::ctbp-1(+)]*). We found that AIA-specific restoration of *ctbp-1* was sufficient to suppress *P_ceh-28_::gfp* misexpression in an otherwise *ctbp-1* mutant background (Figs. 1F; S2A), demonstrating that *ctbp-1* is able to act cell-autonomously to regulate *ceh-28* expression in the AIA neurons.

To determine if *ctbp-1* can act in older animals to suppress AIA gene misexpression, we generated a transgenic construct that drives expression of wild-type *ctbp-1* throughout the worm in response to a short heat shock, *nEx2351[P_hsp-16.2_::ctbp-1(+);P_hsp-16.41_::ctbp-1(+)]* (hereafter referred to as *nEx2351[P_hsp_::ctbp-1(+)]*). We found that heat shock during the L4 larval stage was sufficient to suppress *P_ceh-28_::gfp* misexpression in adult *ctbp-1* mutant AIAs (Figs. 1G-H; S2B), demonstrating that *ctbp-1* can act in L4-to-young adult stage worms to regulate AIA gene expression.

From these data we conclude that *ctbp-1* is able to act cell-autonomously and in L4-to-young adult worms to prevent expression of at least one non-AIA gene in the AIA neurons.

### *ctbp-1* mutant AIAs are not transdifferentiating into an M4-like cell identity

We asked if *P_ceh-28_::gfp* misexpression in the AIAs of *ctbp-1* mutants might be a consequence of the AIAs transdifferentiating into an M4-like cell identity. We scored *ctbp-1* mutants for cell-type markers expressed in, although not necessarily unique to, either M4 or the AIA neurons (Figs. 2A-B; S3A-B). We found that *ctbp-1* mutant AIAs expressed all five of five AIA markers tested and did not express any of four other (non-*ceh-28)* M4 markers tested. Of particular note, *ctbp-1* mutant AIAs did not misexpress either of the two tested M4 genes known to be directly regulated by *ceh-28* (i.e. *dbl-1* and *egl-17)*, indicating that the *ceh-28* misexpression in mutant AIAs does not activate the *ceh-28* regulatory pathway [61, 62]. We conclude that *ctbp-1* mutant AIAs are not transdifferentiated into M4-like cells and instead seem to retain much of their AIA identity while gaining at least one M4 characteristic (i.e. *ceh-28* expression) later in life.

**Figure 2.**
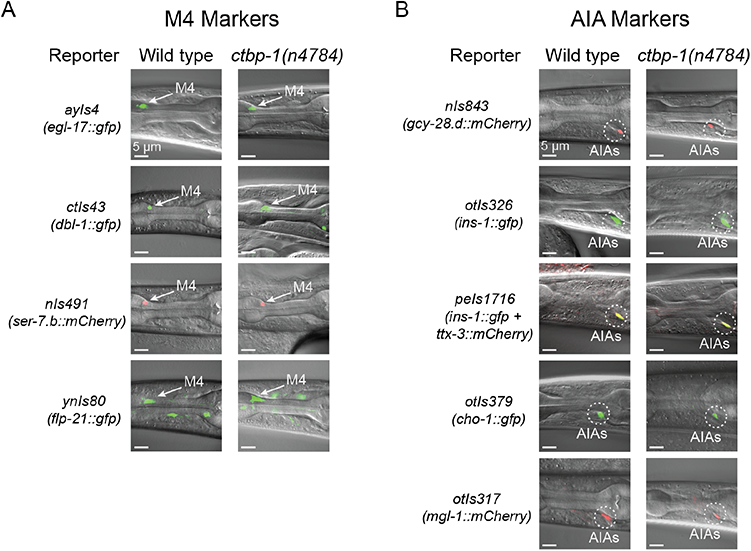
*ctbp-1* mutant AIAs retain multiple aspects of their AIA gene expression profile. (A-B) Expression of (A) M4 markers *egl-17*, *dbl-1*, *ser-7.b* and *flp-21* and (B) AIA markers *gcy-28.d*, *ins-1*, *ttx-3*, *cho-1* and *mgl-1* in wild-type (left image) and *ctbp-1(n4784)* (right image) L4 larval worms. Arrow, M4 neuron. Circles, AIAs. Scale bar, 5 µm. Images are oriented such that left corresponds to anterior, top to dorsal.

### *ctbp-1* mutants display an increasingly severe disruption of AIA morphology

Because of the time-dependency of the defect of *ctbp-1* mutants in AIA cell identity, we hypothesized that *ctbp-1* might act to maintain the AIA cell identity. To test this hypothesis, we examined morphological and functional aspects of AIA identity at both early and late larval stages. To assay AIA morphology, we generated a transgenic construct driving expression of GFP throughout the AIA cell (*nIs840[P_gcy-28.d_::gfp]*). We crossed this construct into *ctbp-1* mutant worms and visualized AIA morphology in L1 and L4 larvae as well as in day 1 adults (Fig. 3A). We found that L1 *ctbp-1* mutant AIAs appeared grossly wild-type in morphology (Fig. 3A). However, L4 and adult *ctbp-1* mutant AIAs had ectopic neurite branches that extended from both the anterior and posterior ends of the AIA cell body (Fig. 3A). The penetrance of these ectopic branches increased progressively in later larval stage and adult mutants (Fig. 3B-C). Older *ctbp-1* mutant AIAs also appeared to have an elongated cell body compared to wild-type AIAs. Quantification of this defect revealed that L4 and adult mutant AIA cell bodies, but not those of L1s, were significantly longer than their wild-type counterparts (Fig. 3D). To assess if this increase in AIA length was a consequence of an increase in AIA size, we measured the maximum area of the AIA cell body from cross-sections of these cells. We found that the maximum area of the AIA cell body did not significantly differ between wild-type and mutant AIAs at any stage (Fig. S4A), indicating that mutant AIAs were misshapen but not enlarged. To confirm that we were not biased by an awareness of genotype while measuring AIA lengths, we blinded the wild-type and *ctbp-1* AIA images used for length measurements and scored the blinded images as either “normal” or “elongated” (Fig. S4B). Again, at the L1 larval stage both wild-type and *ctbp-1* mutant AIAs appeared overwhelmingly “normal,” whereas at both the L4 larval stage and in day 1 adults *ctbp-1* mutant AIAs were scored as “elongated” at a consistently higher rate than their wild-type counterparts. Collectively, these results demonstrate that *ctbp-1* mutant AIAs display abnormal morphology and that the severity of the observed morphological defects in *ctbp-1* mutants increases from L1 to L4 to adulthood. Furthermore, the relative lack of AIA morphological defects in L1 *ctbp-1* mutants suggests that *ctbp-1* is not required for the establishment of proper AIA morphology but instead acts to maintain AIA morphology over time.

**Figure 3.**
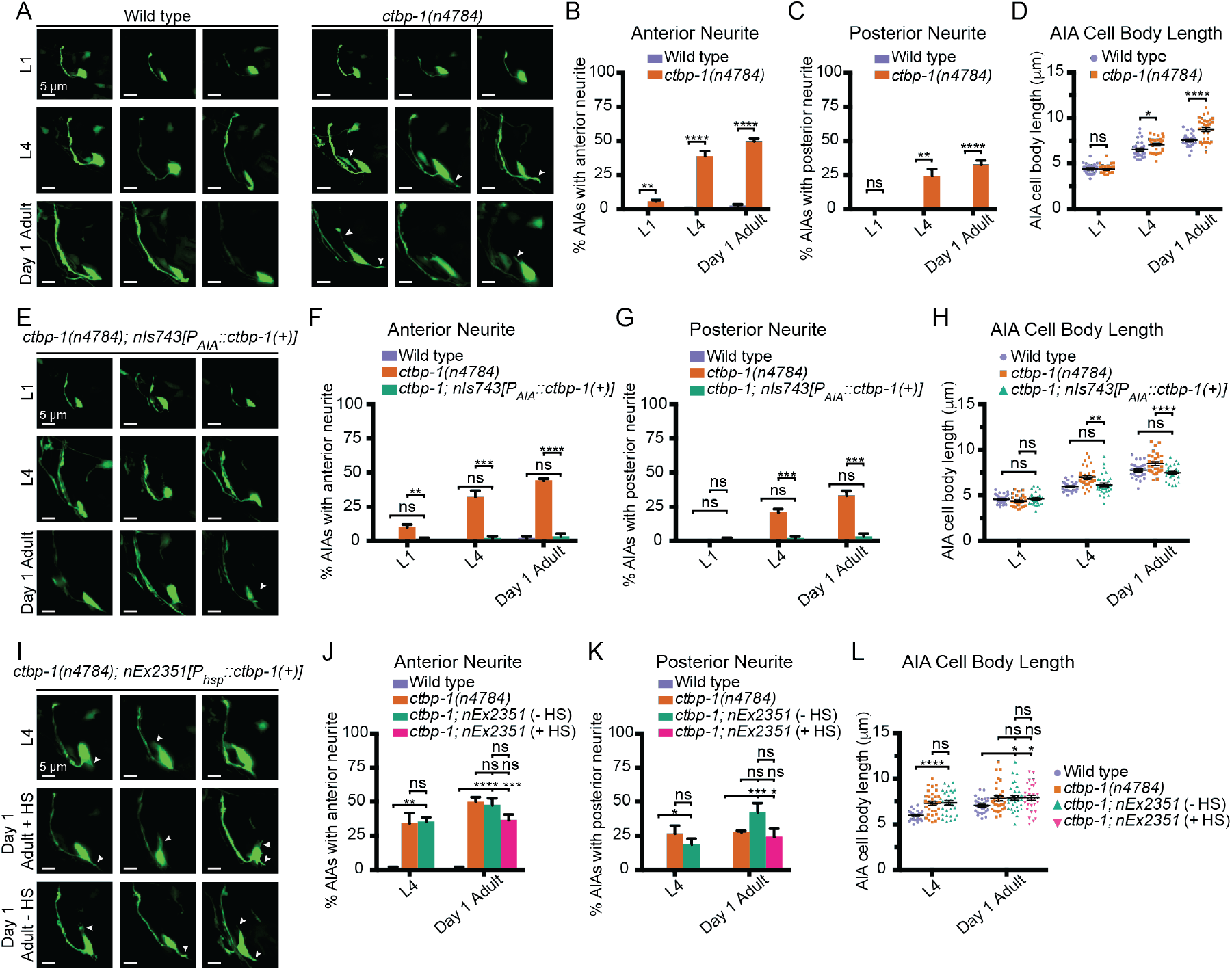
Loss of *ctbp-1* results in a progressive decline in AIA morphology. (A) Three representative images of an AIA neuron in wild-type (left) and *ctbp-1(n4784)* (right) worms at L1 (top), L4 (middle) and day 1 adult (bottom) stages. Arrows, examples of ectopic neurites protruding from the AIA cell body. Scale bar, 5 µm. (B-C) Percentage of AIAs in wild-type and *ctbp-1* worms at the L1, L4 and day 1 adult stages with an ectopic neurite protruding from the (B) anterior or (C) posterior of the AIA cell body. Mean ± SEM. *n* = 60 AIAs scored per strain per stage, 4 biological replicates. ns, not significant, **p<0.01, ****p<0.0001, unpaired t-test. (D) Quantification of AIA cell body length in wild-type and *ctbp-1* worms at the L1, L4 and day 1 adult stages. Mean ± SEM. *n* = 30 AIAs scored per strain per stage. ns, not significant, *p<0.05, ****p<0.0001, unpaired t-test. (E) Three representative images of an AIA neuron in *ctbp-1; nIs743[P_gcy-28.d_::ctbp-1(+)]* worms at L1 (top), L4 (middle) and day 1 adult (bottom) stages. Arrows, examples of ectopic neurites protruding from the AIA cell body. Scale bar, 5 µm. (F-G) Percentage of AIAs in wild-type, *ctbp-1* and *ctbp-1; nIs743* worms at the L1, L4 and day 1 adult stages with an ectopic neurite protruding from the (F) anterior or (G) posterior of the AIA cell body. Mean ± SEM. *n* = 30 AIAs scored per strain per stage, 3 biological replicates. ns, not significant, **p<0.01, ***p<0.001, ****p<0.0001, one-way ANOVA with Tukey’s correction. (H) Quantification of AIA cell body length in wild-type, *ctbp-1* and *ctbp-1; nIs743* worms at the L1, L4 and day 1 adult stages. Mean ± SEM. *n* ≥ 30 AIAs scored per strain per stage. ns, not significant, **p<0.01, ****p<0.0001, one-way ANOVA with Tukey’s correction. (I) Three representative images of an AIA neuron in *ctbp-1; nEx2351[P_hsp-16.2_::ctbp-1(+);P_hsp-16.41_::ctbp-1(+)]* worms at L4 (top), day 1 adult with heat shock (HS) (middle) and day 1 adult without heat shock (bottom). Arrows, examples of ectopic neurites protruding from the AIA cell body. Scale bar, 5 µm. (J-K) Percentage of AIAs in wild-type, *ctbp-1* and *ctbp-1; nEx2351* worms at L4 and day 1 adult (with or without heat shock) stages with an ectopic neurite protruding from the (J) anterior or (K) posterior of the AIA cell body. Mean ± SEM. *n* = 30 AIAs scored per strain per stage, 3 biological replicates. ns, not significant, *p<0.05, **p<0.01, ***p<0.001, ****p<0.0001, one-way ANOVA with Tukey’s correction. (L) Quantification of AIA cell body length in wild-type, *ctbp-1* and *ctbp-1; nEx2351* worms at L4 and day 1 adult (with or without heat shock) stages. Mean ± SEM. *n* ≥ 30 AIAs scored per strain per stage. ns, not significant, *p<0.05, ****p<0.0001, one-way ANOVA with Tukey’s correction. The *ctbp-1* allele used for all panels of this figure was *n4784*. All strains contain *nIs840[P_gcy-28.d_::gfp],* and all strains other than “Wild type” contain *nIs348[P_ceh-28_::mCherry]* (not shown in images). Images are oriented such that left corresponds to anterior, top to dorsal.

We next asked if *ctbp-1* acts cell-autonomously and at later stages to regulate AIA morphology as it does for AIA gene expression. We visualized *ctbp-1* mutant AIAs carrying the AIA-specific *ctbp-1(+)* rescue construct *nIs743[P_AIA_::ctbp-1(+)]* (Fig. 3E). We found that AIA-specific restoration of *ctbp-1* in mutant worms rescued all AIA morphological defects to near-wild-type levels at all stages tested, indicating that *ctbp-1* can act cell-autonomously to regulate AIA morphology (Fig. 3F-H). Next, we visualized *ctbp-1* mutant worms carrying the heat shock-inducible *ctbp-1(+)* rescue construct *nEx2351[P_hsp_::ctbp-1(+)]*. We found that heat shock at the L4 stage did not restore *ctbp-1* mutant AIA morphology in day 1 adults back to wild-type, nor was there any significant difference in the frequency of morphological defects between heat-shocked and non-heat-shocked adults carrying the rescue construct (Fig. 3I-L). We speculate that the lack of restoration of morphology in late-stage worms might be a consequence of the defects being irreversible, and that *ctbp-1* might be continuously required to prevent such defects from occurring. From these data we conclude that *ctbp-1* can act cell-autonomously, and possibly continuously, to maintain aspects of AIA morphology in a manner similar to AIA gene expression.

### *ctbp-1* mutants display a progressive decline of AIA function

The AIA interneurons integrate sensory information from a number of sensory neurons, resulting in modulation of the movement of the worm in response to environmental stimuli [63–65]. The AIAs function in response to volatile odors and play an important role in learning associated with the sensation of volatile odors or salts [63, 66]. We asked if *ctbp-1* mutants are abnormal in a behavior known to require the AIAs – adaptation to the volatile odor 2-butanone [66] – reasoning that if AIA function is disrupted in *ctbp-1* mutants, there should be a reduction of adaptation (and thus greater attraction) to butanone in *ctbp-1* worms relative to wild-type worms.

Consistent with previous studies [66], we found that worms that had been briefly starved with 90 minutes of food deprivation and had no prior experience with butanone (so-called “naïve” worms) were generally attracted to the odor, while worms that were briefly starved in the presence of butanone (“conditioned” worms) adapted to the odor and exhibited mild repulsion to it (Fig. 4A-E). We next compared wild-type and *ctbp-1* mutant worms for their ability to adapt to butanone. We found that while L1 *ctbp-1* worms showed an ability to adapt to butanone roughly similar to that of their wild-type counterparts, conditioned L4 *ctbp-1* mutants displayed a significant decrease in repulsion from butanone relative to wild-type L4 animals, indicating a decrease in their ability to adapt to the odor (Fig. 4B-E). As a control, we assayed a strain carrying a transgenic construct that genetically ablates the AIA neurons, JN580. As expected, JN580 worms displayed decreased butanone adaptation at both the L1 and L4 larval stages. Thus, *ctbp-1* mutant worms displayed a defect in butanone adaptation similar to that of an AIA-ablated strain and did so only at a later larval stage, suggesting a potential loss of AIA function in L4 *ctbp-1* mutants. However, while *ctbp-1* mutant L4s exhibited weaker butanone adaptation than their wild-type counterparts, this defect was not as severe as that of JN580 L4s, indicating that *ctbp-1* mutant AIAs might retain some function. Additionally, the lack of a butanone adaptation defect in L1 *ctbp-1* mutants similar to that of L1 JN580 worms further suggests that loss of *ctbp-1* does not disrupt early AIA function and shows that *ctbp-1* is not required for the establishment of functional AIA neurons.

**Figure 4.**
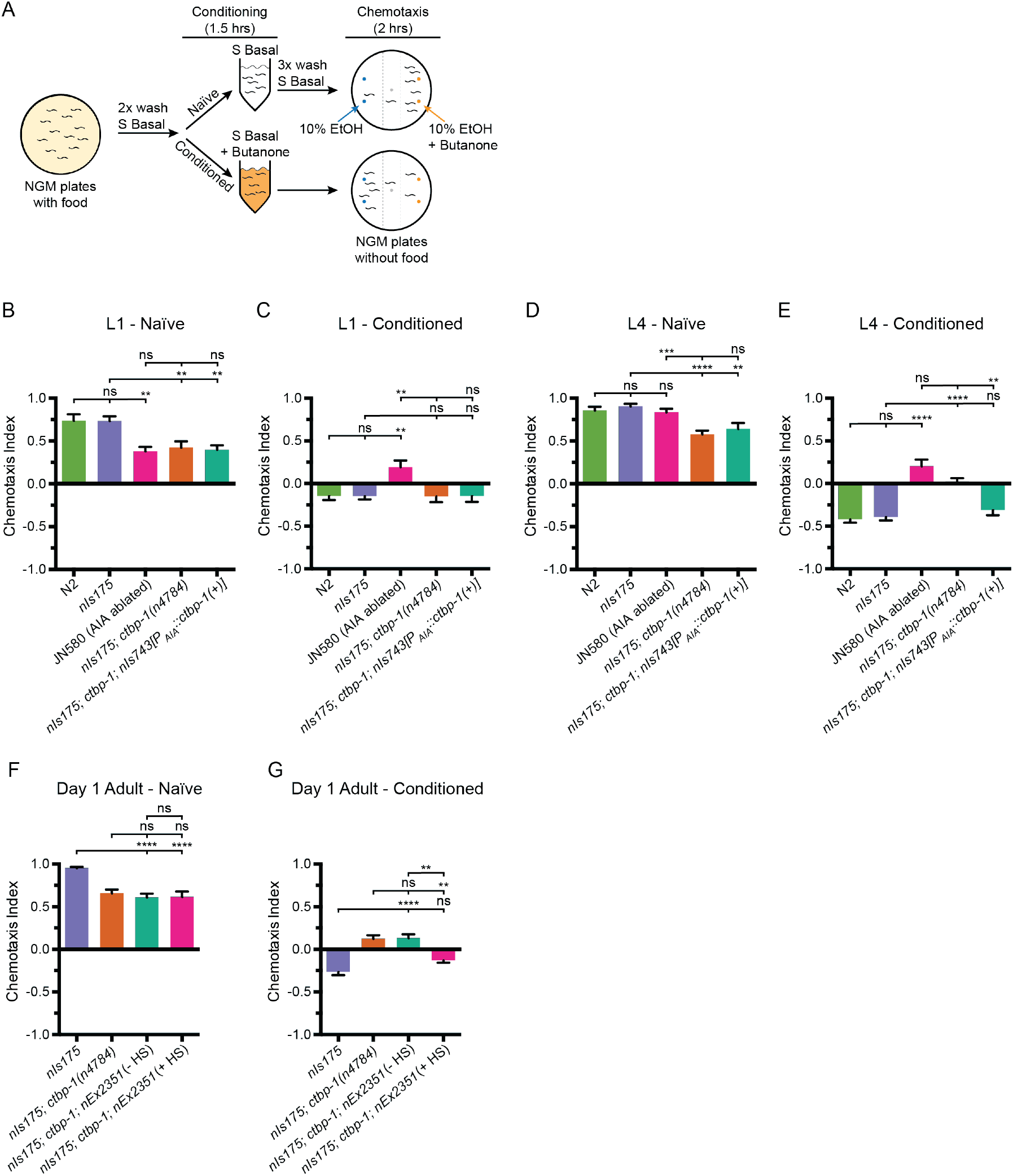
Loss of *ctbp-1* results in a disruption of AIA function in L4-to-day 1 adult worms. (A) Schematic of the butanone adaptation assay. L1 or L4 worms from synchronized populations were washed off plates with S Basal, washed with S Basal, split into naïve and conditioned populations, incubated in S Basal with or without 2-butanone for 1.5 hours, washed again with S Basal, allowed to chemotax for two hours on unseeded plates containing two 1 µl spots of 10% ethanol (blue dots) and 2-butanone diluted in 10% ethanol (orange dots), and then scored. (B-E) Chemotaxis indices of (B,D) naïve or (C,E) conditioned wild-type (N2 and *nIs175*), AIA-ablated (JN580), *nIs175*; *ctbp-1(n4784)*, and *nIs175*; *ctbp-1* mutants containing a transgene driving expression of wild-type *ctbp-1* under an AIA-specific promoter (*nIs743[P_gcy-28.d_::ctbp-1(+)]*) at the (B-C) L1 or (D-E) L4 larval stage. Mean ± SEM. *n* ≥ 6 assays per condition, ≥ 50 worms per assay. ns, not significant, **p<0.01, ***p<0.001, ****p<0.0001, one-way ANOVA with Tukey’s correction. (F-G) Chemotaxis indices of (F) naïve or (G) conditioned *nIs175*, *nIs175*; *ctbp-1*, and *nIs175*; *ctbp-1* mutants carrying the heat-shock-inducible transgene *nEx2351[P_hsp-16.2_::ctbp-1(+);P_hsp-16.41_::ctbp-1(+)]* with or without heat shock (HS) at the day 1 adult stage. Mean ± SEM. *n* ≥ 5 assays per condition, ≥ 50 worms per assay. ns, not significant, **p<0.01, ****p<0.0001, one-way ANOVA with Tukey’s correction. The *ctbp-1* allele used for all panels of this figure was *n4784*.

We next asked if *ctbp-1* can act cell-autonomously in the AIAs and in older worms to regulate butanone adaptation. We assayed *ctbp-1* mutants carrying the AIA-specific rescue construct *nIs743[P_AIA_::ctbp-1(+)]* for butanone adaptation (Fig. 4B-E) and found that AIA-specific restoration of *ctbp-1* rescued butanone adaption of conditioned *ctbp-1* mutant L4s to near wild-type levels (Fig. 4E). We conclude that the butanone adaptation defect of *ctbp-1* mutants is caused by a disruption of AIA function and that *ctbp-1* can act cell-autonomously to regulate this AIA function. Next, we assayed *ctbp-1* mutants carrying the heat shock-inducible *ctbp-1(+)* rescue construct *nEx2351[P_hsp_::ctbp-1(+)]* for butanone adaptation. We found that restoration of *ctbp-1* by heat shock at the L4 larval stage rescued the butanone adaptation defect in day 1 adults, indicating that *ctbp-1* can act in L4-to-day 1 adult worms to maintain proper AIA function after the initial establishment of the AIA cell identity (Fig. 4F-G). Taken together, these data establish that loss of *ctbp-1* disrupts the function of the AIA neurons and that *ctbp-1* can act cell-autonomously and in L4-to-day 1 adult worms to maintain AIA function.

While conducting these assays, we observed that naïve *ctbp-1* mutant worms displayed a mildly weaker attraction to butanone than did their wild-type counterparts at both the L1 and L4 larval stages (Fig. 4B,D). AIA-specific rescue of *ctbp-1* did not rescue this mild chemotaxis defect – naïve *ctbp-1* mutants carrying the *P_AIA_::ctbp-1(+)* construct still displayed weaker butanone attraction than wild-type worms (Fig. 4B,D). We suggest that this defect in attraction to butanone is not a consequence of dysfunction of the AIAs but rather of some other cell(s) involved in butanone chemotaxis. Consistent with this hypothesis, we found that *ctbp-1* mutants were defective in chemotaxis to the volatile odor isoamyl alcohol but were not defective in the response to diacetyl (Fig. S5A-B) while AIA-ablated strain JN580 animals were not defective in either response (Fig. S5A-B). These observations indicate that *ctbp-1* mutant worms have a broader defect in chemotaxis caused by the disruption of the function of cells other than the AIAs. Because our primary focus has been on how *ctbp-1* functions to maintaining the AIA cell identity, we did not attempt to identify the other cells with functions perturbed by the loss of *ctbp-1*.

### *ctbp-1* mutant AIAs have additional defects in gene expression

To better characterize the genetic changes occurring in mutant AIAs, we performed a single, exploratory single-cell RNA-Sequencing (scRNA-Seq) experiment comparing wild-type and *ctbp-1* mutant worms. We sequenced RNA from the neurons of wild-type and *ctbp-1* L4 worms and processed the resulting data using the 10X CellRanger pipeline to identify presumptive AIA neurons based on the expression of several AIA markers (*gcy-28*, *ins-1*, *cho-1*) shown above to be expressed in both wild-type and *ctbp-1* mutant AIAs (Fig. 2B). Confirming that these data captured changes in the AIA transcriptional profiles, we found that *ctbp-1* mutant AIAs showed high levels of expression of *ceh-28,* while wild-type AIAs showed no detectable *ceh-28* expression (Fig. S6).

We analyzed AIA transcriptional profiles to identify genes that appeared to be either expressed in *ctbp-1* mutant AIAs and not expressed in wild-type AIAs (similar to *ceh-28*) or expressed in wild-type AIAs but not expressed in *ctbp-1* AIAs. To confirm candidate genes, we crossed existing reporters for those genes to *ctbp-1* mutants or, in cases for which reporters were not readily available, generated our own transgenic constructs. We identified and confirmed one gene that, similar to *ceh-28*, was not expressed in wild-type AIAs but was misexpressed in *ctbp-1* mutant AIAs: *acbp-6*, which is predicted to encode an acyl-Coenzyme A binding protein [67] (Fig. 5A; S6). We also identified and confirmed two genes expressed in wild-type AIAs but not expressed in *ctbp-1* mutant AIAs: *sra-11*, which encodes a transmembrane serpentine receptor [68]; and *glr-2,* which encodes a glutamate receptor [69] (Fig. 5C,E; S6). We visualized the *acbp-6* reporter *nEx3081[P_acbp-6_::gfp]*, the *sra-11* reporter *otIs123[P_sra-11_::gfp]* and the *glr-2* reporter *ivEx138[P_glr-2_::gfp]* in wild-type and *ctbp-1* L4 worms and confirmed that *acbp-6* was absent in wild-type AIAs but misexpressed in *ctbp-1* mutants (Fig. 5A-B), while both *sra-11* and *glr-2* were consistently expressed in wild-type AIAs but not expressed in the AIAs of *ctbp-1* mutants (Fig. 5C-F). We also visualized these reporters in L1 wild-type and *ctbp-1* worms and found that both *P_acbp-6_::gfp* and *P_sra-11_::gfp* displayed a time-dependence to their expression similar to that of *P_ceh-28_::gfp* − *P_acbp-6_::gfp* was rarely detectible in the AIAs of either wild-type of *ctbp-1* AIAs at the L1 stage but was consistently expressed in *ctbp-1* mutant L4 AIAs (Fig. 5A-B), while *P_sra-11_::gfp* was rarely detectible in the AIAs of either wild-type or *ctbp-1* mutant L1 worms but was expressed in the AIAs of most wild-type worms by the L4 stage while remaining off in the AIAs of most L4 *ctbp-1* mutants (Fig. 5C-D). These observations suggest that, like *ceh-28* expression, *acbp-6* and *sra-11* expression is regulated by *ctbp-1* primarily in the AIAs of late-stage larvae and adults. By contrast, *glr-2* was expressed in wild-type but not *ctbp-1* AIAs in both L1 and L4 larvae (Fig. 5E-F).

**Figure 5.**
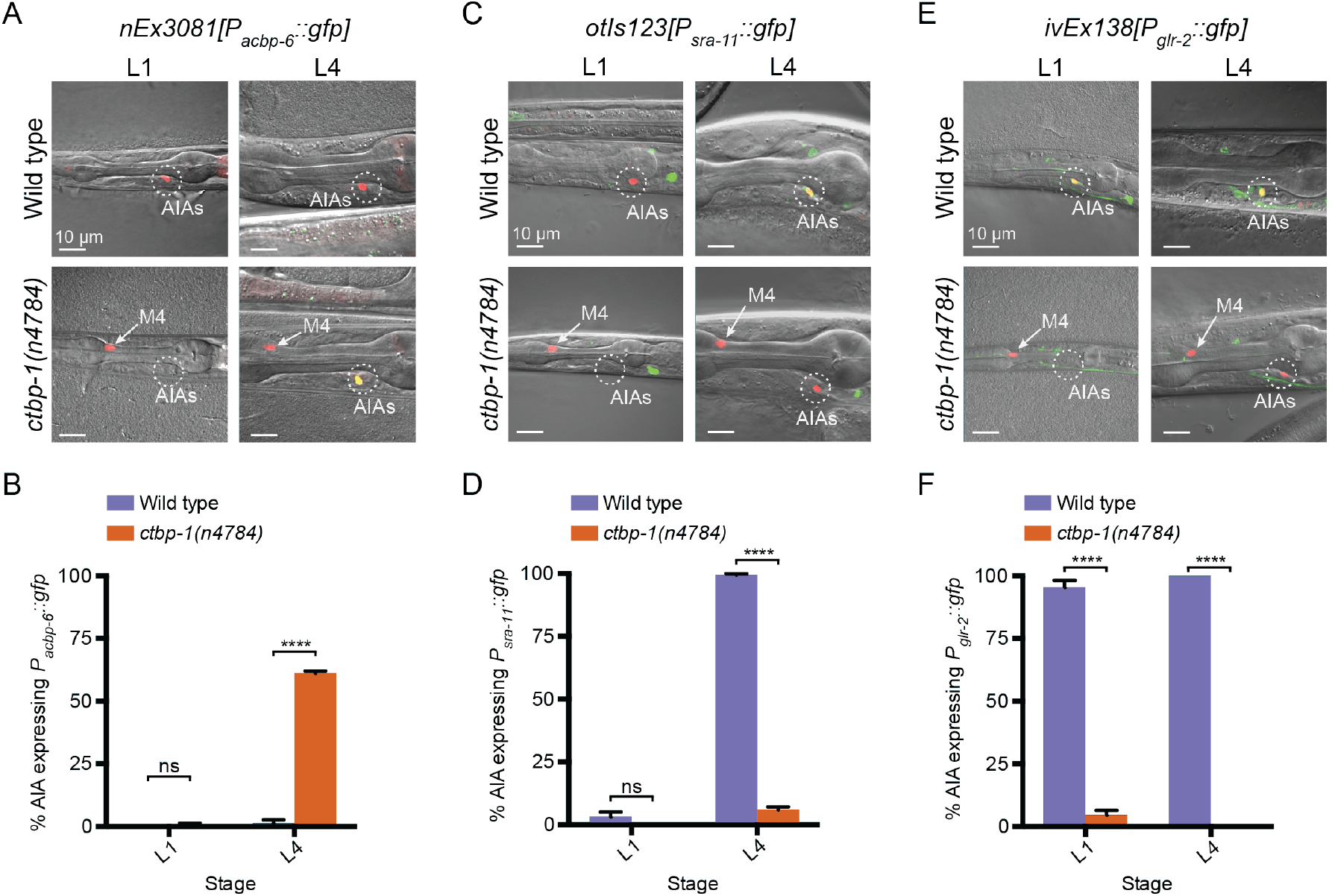
Loss of *ctbp-1* results in a disruption to normal AIA gene expression. (A,C,E) (A) *nEx3081[P_acbp-6_::gfp]*, (C) *otIs123[P_sra-11_::gfp],* or (E) *ivEx138[P_glr-2_::gfp]* expression in wild-type (top) and *ctbp-1(n4784)* (bottom) worms at the L1 larval stage (left) and L4 larval stage (right). Wild-type strains contain *nIs843[P_gcy-28.d_::mCherry]. ctbp-1* mutant strains contain *nIs348[P_ceh-28_::mCherry].* Arrow, M4 neuron. Circle, AIAs. Scale bar, 10 µm. (B,D,F) Percentage of wild-type and *ctbp-1(n4784)* expressing (B) *P_acbp-6_::gfp*, (D) *P_sra-11_::gfp,* or (F) *P_glr-2_::gfp* in the AIA neurons at L1 and L4 larval stages. Wild-type strains contain *nIs843[P_gcy-28.d_::mCherry]. ctbp-1* mutant strains contain *nIs348[P_ceh-28_::mCherry].* Mean ± SEM. *n* ≥ 50 worms per strain per stage, 3 biological replicates. ns, not significant, ****p<0.0001, unpaired t-test.

These data demonstrate that mutant AIAs fail to turn on and/or maintain the expression of genes characteristic of the adult AIA neuron (*sra-11* and *glr-2*) while misexpressing at least two genes uncharacteristic of AIA (*ceh-28* and *acbp-6)*. That the majority of these abnormalities in AIA gene expression occurred long after the AIAs are generated during embryogenesis further supports the conclusion that *ctbp-1* does not act to establish the AIA cell identity.

Collectively, our findings concerning AIA gene expression, morphology and function demonstrate that *ctbp-1* acts to maintain the AIA cell identity, plays little to no role in the initial establishment of the AIA cell fate, and can act cell-autonomously and in older worms to maintain these aspects of the AIA identity.

### *egl-13* mutations suppress the *ctbp-1* mutant phenotype

To investigate how *ctbp-1* acts to maintain AIA cell identity, we performed a mutagenesis screen for suppression of *P_ceh-28_::gfp* misexpression in the AIAs of L4 *ctbp-1* mutants (Fig. 6A). Using a combination of Hawaiian SNP mapping [70] and whole-genome sequencing, we identified the gene *egl-13*, which encodes a SOX family transcription factor, as a suppressor of *ctbp-1*. *egl-13* has been shown to act in the establishment of the BAG and URX cell fates and in vulval development of *C. elegans* [56, 71], and its mammalian orthologs SOX5 and SOX6 act in neural fate determination [72, 73]. We isolated three alleles of *egl-13* as *ctbp-1* suppressors: *n5937*, a mutation of the splice acceptor site at the beginning of the 6^th^ exon of the *egl-13a* isoform resulting in a frameshift and early stop; *n6013*, a Q381ochre nonsense mutation towards the end of the *egl-13* transcript; and *n6313*, a 436-nucleotide deletion spanning the 7^th^ and 8^th^ exons of the *egl-13a* isoform (Figs. 6B; S7A-B). We generated and introduced a transgenic construct carrying a wild-type copy of *egl-13* under its native promoter into these mutant strains and found that this construct was capable of rescuing the suppression of *P_ceh-28_::gfp* misexpression by all three *egl-13* alleles, demonstrating that loss of *egl-13* function suppresses this aspect of the *ctbp-1* mutant phenotype and suggesting that these alleles are likely loss-of-function alleles of *egl-13* (Fig. S7C).

**Figure 6.**
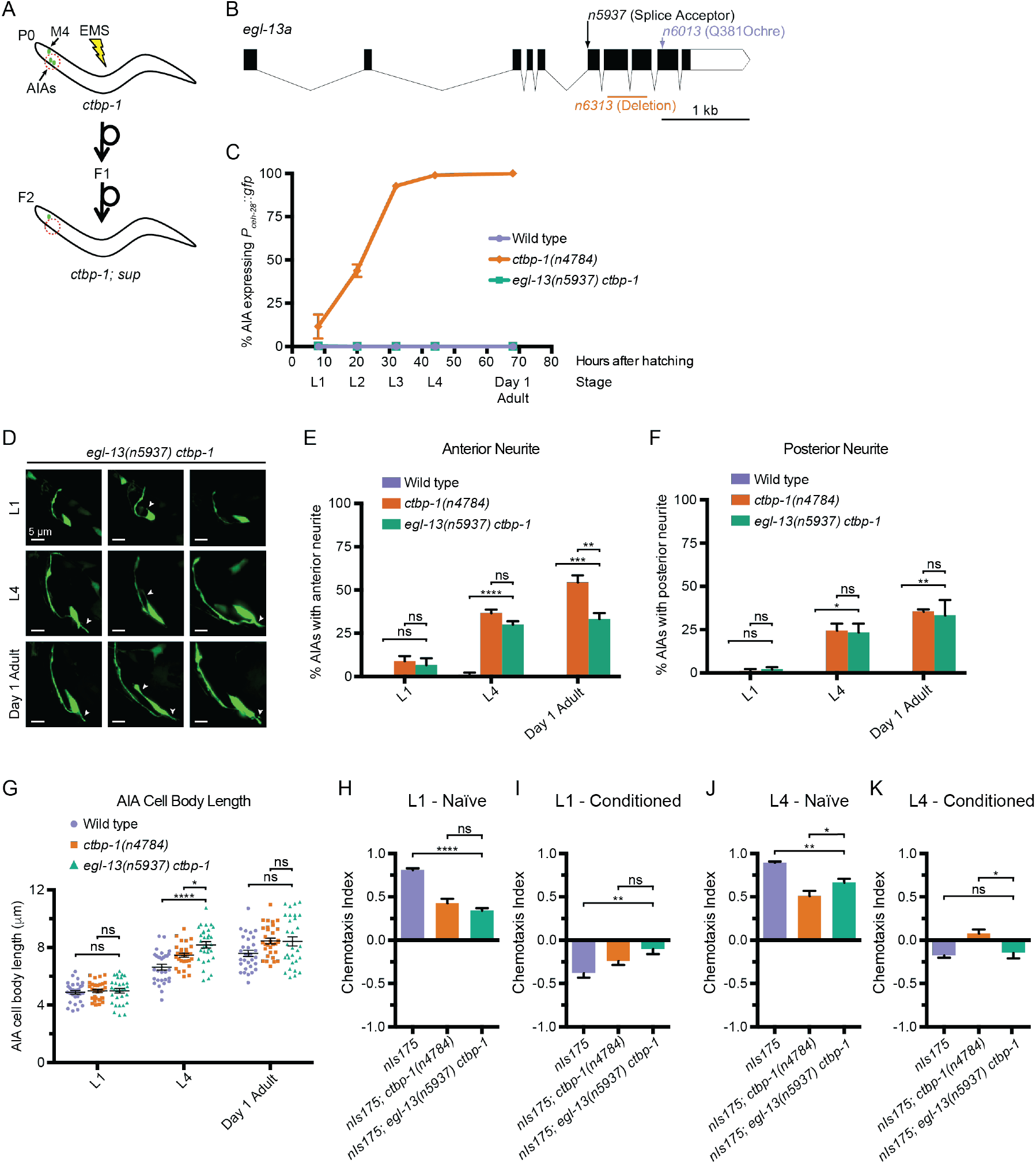
A suppressor screen reveals *egl-13* as a *ctbp-1* genetic interactor. (A) Schematic of *ctbp-1* suppressor screen design. *ctbp-1* mutant worms carrying *nIs175[P_ceh-28_::gfp]* were mutagenized with ethyl methanesulfonate (EMS), and their F2 progeny were screened for continued *nIs175* expression in M4 and loss of expression in the AIA neurons (red circle). (B) Gene diagram of the *egl-13a* isoform. Arrows (above), point mutations. Line (below), deletion. Scale bar (bottom right), 1 kb. (C) Percentage of wild-type, *ctbp-1* and *egl-13(n5937)* worms expressing *nIs175* in the AIA neurons over time. Time points correspond to the L1, L2, L3, L4 larval stages, and day 1 adult worms (indicated below X axis). All strains contain *nIs175[P_ceh-28_::gfp].* Mean ± SEM. *n* ≥ 100 worms per strain per stage, 3 biological replicates. (D) Three representative images of an AIA neuron in *egl-13 ctbp-1* worms at L1 (top), L4 (middle) and day 1 adult (bottom) stages. Arrows, examples of ectopic neurites protruding from the AIA cell body. Image oriented such that left corresponds to anterior, top to dorsal. Scale bar, 5 µm. (E-F) Percentage of AIAs in wild-type, *ctbp-1* and *egl-13 ctbp-1* worms at the L1, L4 and day 1 adult stages with an ectopic neurite protruding from the (E) anterior or (F) posterior of the AIA cell body. Mean ± SEM. *n* = 30 AIAs scored per strain per stage, 3 biological replicates. ns, not significant, *p<0.05, **p<0.01, ***p<0.001, ****p<0.0001, one-way ANOVA with Tukey’s correction. (G) Quantification of AIA cell body length in wild-type, *ctbp-1* and *egl-13 ctbp-1* worms at the L1, L4 and day 1 adult stages. Mean ± SEM. *n* ≥ 30 AIAs scored per strain per stage. ns, not significant, *p<0.05, ****p<0.0001, one-way ANOVA with Tukey’s correction. (H-K) Chemotaxis indices of (H,J) naïve or (I,K) conditioned wild-type, *ctbp-1* and *egl-13 ctbp-1* worms at the (H-I) L1 or (J-K) L4 larval stage. Mean ± SEM. *n* ≥ 5 assays per condition, ≥ 50 worms per assay. ns, not significant, *p<0.05, **p<0.01, ****p<0.0001, one-way ANOVA with Tukey’s correction. The *ctbp-1* allele used for all panels of this figure was *n4784*. The *egl-13* allele used for all panels of this figure was *n5937*. All strains in (D-G) contain *nIs840[P_gcy-28.d_::gfp]* and all strains in (D-G) other than “Wild type” contain *nIs348[P_ceh-28_::mCherry]* (not shown in images). All strains in (C, H-K) contain *nIs175[P_ceh-28_::gfp]*.

We assayed the loss-of-function allele of *egl-13* with the highest penetrance of suppression*, n5937,* for its ability to suppress *P_ceh-28_::gfp* misexpression over the course of larval development and into adulthood of *ctbp-1* mutant worms (Fig. 6C). *egl-13(n5937)* strongly suppressed *ctbp-1* at all stages, resulting in little to no misexpression of *P_ceh-28_::gfp* in the AIAs of *egl-13 ctbp-1* double mutants at any larval stage or in day 1 adults.

To determine if, like *ctbp-1*, *egl-13* can act cell-autonomously in the AIAs, we generated a transgenic construct that drives expression of a wild-type copy of *egl-13* in the AIAs (*nEx3055[P_gcy-28.d_::egl-13(+)]*). Introduction of this construct to *egl-13(n5937) ctbp-1* double mutants rescued the *egl-13* suppression of *P_ceh-28_::gfp* misexpression in the AIAs, indicating that *egl-13* can function cell-autonomously (Fig. S8A-B). These results suggest that, in the absence of *ctbp-1* function, ectopic *egl-13* activity drives *ceh-28* misexpression, and thus that *ctbp-1* likely normally acts to repress *egl-13* activity in the AIAs.

We next asked if *egl-13(n5937)* could suppress the AIA morphological and functional defects of *ctbp-1* mutants. To both test suppression of AIA morphological defects and confirm the presence of the AIA neurons in *egl-13 ctbp-1* double mutants, we crossed the AIA morphology reporter *nIs840[P_gcy-28.d_::gfp]* into *egl-13 ctbp-1* double mutants and scored AIA morphology in L1, L4 and day 1 adult worms (Fig. 6D-G). *egl-13 ctbp-1* double mutant AIAs displayed a mild (though significant) reduction in the penetrance of ectopic anterior neurites only in adult worms, no significant change in the frequency of posterior neurites at any stage, and a slight increase in AIA cell body length of L4 worms (though the difference was no longer significant in adults). These data demonstrate that loss of *egl-13* has little consistent effect on the AIA morphological defects caused by a loss of *ctbp-1* activity, suggesting that *ctbp-1* maintains AIA morphology primarily through *egl-13*-independent pathways.

We next assayed the ability of *egl-13(n5937)* to suppress AIA functional defects. We tested *egl-13 ctbp-1* double mutants for butanone adaptation and found that, at the L1 larval stage, this double mutant strain displayed a detectable response to butanone similar to *ctbp-1* single mutants (Fig. 6H-I). By contrast, at the L4 larval stage, mutation of *egl-13* strongly suppressed the *ctbp-1* mutant defect in butanone adaptation, causing near wild-type levels of repulsion in conditioned worms (Fig. 6J-K). These results indicate that loss of *egl-13* activity suppressed AIA functional defects of *ctbp-1* mutant worms and suggest that *ctbp-1* maintains at least this aspect of AIA cellular function primarily through an *egl-13*-dependent pathway.

From these data we conclude that *ctbp-1* maintains AIA function and at least some aspects of AIA gene expression by antagonizing *egl-13* function and that *ctbp-1* likely also acts primarily through one or more *egl-13-*independent pathways to maintain AIA cellular morphology.

### *egl-13* regulates AIA function through control of *ceh-28* expression

We next asked if mutation of *egl-13* could suppress other *ctbp-1* mutant AIA gene expression defects besides that of *ceh-28*. We crossed in *acbp-6*, *sra-11* and *glr-2* reporters to *egl-13 ctbp-1* double mutants and visualized reporter expression at the L1 and L4 larval stages. We found that mutation of *egl-13* suppressed *P_acbp-6_::gfp* misexpression in the AIAs (Fig. 7A,D, compare to Fig. 5), just as *egl-13* mutation suppressed *P_ceh-28_::gfp* misexpression. By contrast, mutation of *egl-13* had no effect on the loss of *P_sra-11_::gfp* or *P_glr-2_::gfp* expression in *ctbp-1* mutants (Fig. 7B-D). We speculated that EGL-13 might directly regulate expression of *ceh-28* or *acbp-6*. To test this hypothesis, we examined the *ceh-28* and *acbp-6* promoter regions for possible EGL-13 binding sites. We failed to identify any promising candidates, suggesting that regulation of these genes by EGL-13 is likely indirect. These results demonstrate that some, though not all, of the AIA gene expression defects seen in *ctbp-1* mutants are regulated through *egl-13*.

**Figure 7.**
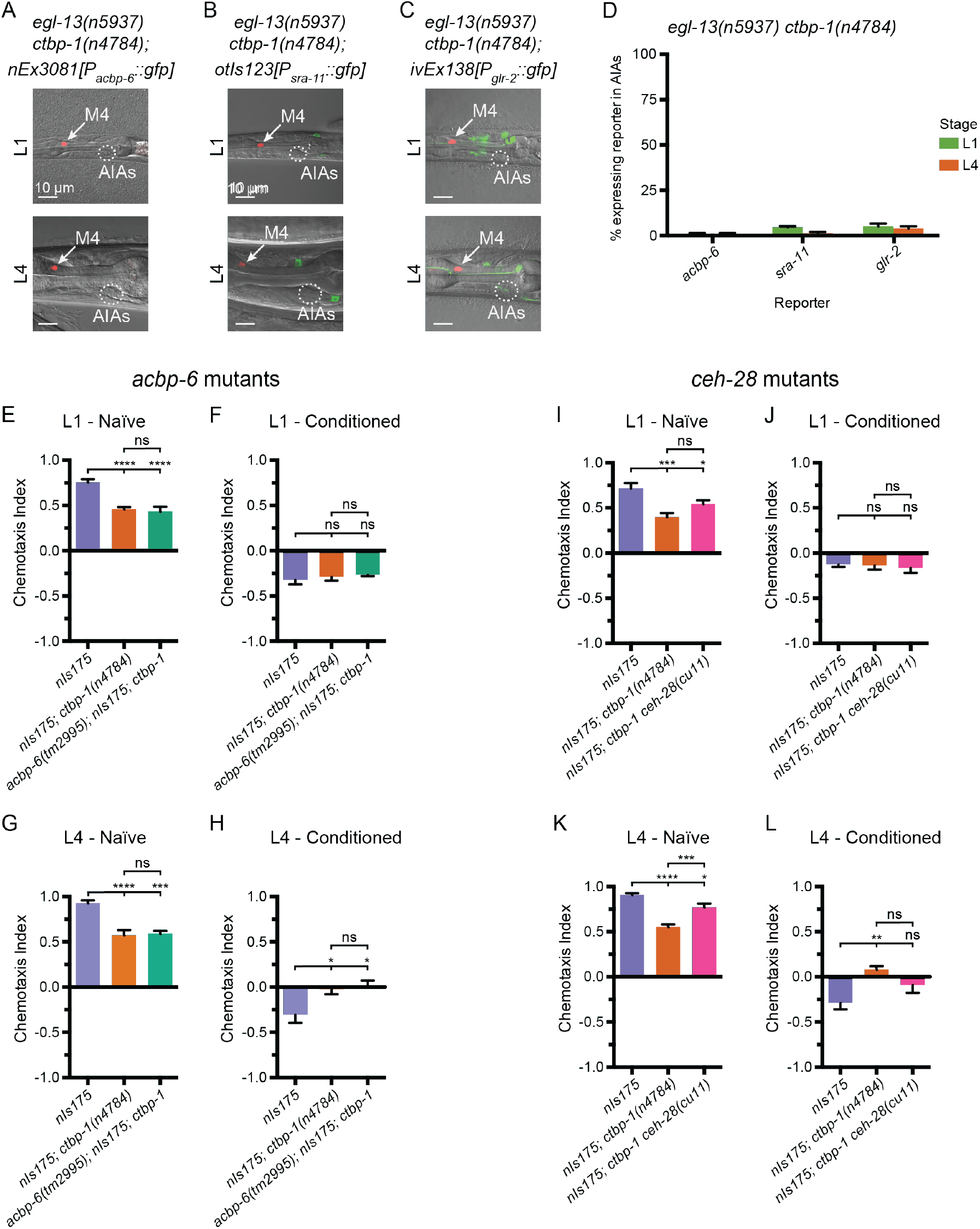
EGL-13 disrupts AIA function partially through driving misexpression of *ceh-28* in *ctbp-1* mutants. (A-C) Expression of markers for AIA misexpressed genes (A) *nEx3081[P_acbp-6_::gfp],* (B) *otIs123[P_sra-11_::gfp]*, or (C) *ivEx138[P_glr-2_::gfp]* in *egl-13(n5937) ctbp-1(n4784)* double mutants at the (top) L1 and (bottom) L4 larval stages. Arrow, M4 neuron. Circle, AIAs. Scale bar, 10 µm. (D) Percentage of *egl-13(n5937) ctbp-1(n4784)* double mutants expressing the indicated reporter in the AIA neurons at the L1 and L4 larval stages. Mean ± SEM. *n* ≥ 50 worms scored per strain, 3 biological replicates. (E-H) Chemotaxis indices of (E,G) naïve or (F,H) conditioned wild-type (*nIs175*), *nIs175; ctbp-1(n4784)*, and *acbp-6(tm2995)*; *nIs175; ctbp-1* mutants at the (E-F) L1 or (G-H) L4 larval stage. Mean ± SEM. *n* ≥ 6 assays per condition, ≥ 50 worms per assay. ns, not significant, *p<0.05, ***p<0.001, ****p<0.0001, one-way ANOVA with Tukey’s correction. (I-L) Chemotaxis indices of (I,K) naïve or (J,L) conditioned wild-type (*nIs175*), *nIs175; ctbp-1(n4784)*, and *nIs175; ctbp-1 ceh-28(cu11)* mutants at the (I-J) L1 or (K-L) L4 larval stage. Mean ± SEM. *n* ≥ 6 assays per condition, ≥ 50 worms per assay. ns, not significant, *p<0.05, ***p<0.001, one-way ANOVA with Tukey’s correction. The *ctbp-1* allele used for all panels of this figure was *n4784*. All strains in Fig. 7A-D contain *nIs348[P_ceh-28_::mCherry]*. Images are oriented such that left corresponds to anterior, top to dorsal.

As *egl-13* is required for misexpression of both *ceh-28* and *acbp-6* as well as for disruption of AIA function in *ctbp-1* mutants, we hypothesized that misexpressed *ceh-28* or *acbp-6* might be contributing to the observed AIA functional defect in *ctbp-1* mutants. If so, we expected that mutations that eliminated the functions of these ectopically expressed genes should restore AIA function in *ctbp-1* mutants. To test this hypothesis, we crossed mutant alleles of *ceh-28 (cu11)* or *acbp-6 (tm2995)* (both deletion alleles spanning greater than half their respective genes) to *ctbp-1(n4784)* mutants and assayed the resulting double mutants for butanone adaptation in L1 and L4 worms. We found that *acbp-6; ctbp-1* double mutants were nearly identical to both naïve and conditioned *ctbp-1* single mutants at both the L1 and L4 larval stages (Fig. 7E-H), indicating that misexpressed *acbp-6* is likely not responsible for the observed AIA functional defect. Conditioned *ctbp-1 ceh-28* double mutants appeared similar to both the wild type and *ctbp-1* single mutants at the L1 stage (Fig. 7I-J). These double mutants display an minor, though not statistically significant, difference in adaptation at the L4 larval stage when compared to either wild-type or *ctbp-1* animals (Fig. 7K-L). We speculate that overexpression of *ceh-28,* caused by a loss of *ctbp-1* and likely driven by ectopic *egl-13* activity, might partially account for the defect in butanone adaptation seen in L4 and day 1 adult *ctbp-1* mutants and that removal of *egl-13* in part restores AIA function by eliminating *ceh-28* misexpression. We propose that *ctbp-1* functions to maintain aspects of the AIA cell identity by preventing *egl-13* from promoting *ceh-28* expression and that *ceh-28* misexpression can perturb proper AIA function. These results also indicate that *ceh-28* overexpression alone is not solely responsible for the observed AIA functional defect, suggesting that the regulation of other, as-of-yet unidentified genes controlled by *ctbp-1* (and potentially *egl-13*) also contribute to the maintenance of the AIA cell identity.

## Discussion

We have shown that the *ctbp-1* transcriptional corepressor gene is required to maintain AIA cell identity and that *ctbp-1* negatively and selectively regulates the function of the *egl-13* transcription factor gene. We suggest that the CTBP-1 protein functions as a transcriptional corepressor to selectively regulate the transcriptional output (either directly or indirectly) of the EGL-13 protein. c*tbp-1* mutant AIAs undergo a progressive decline in their initially wild-type gene-expression pattern, morphology and function. *ctbp-1* can act cell-autonomously and is able to act in older animals to maintain these aspects of the AIA identity. We conclude that CTBP-1 functions to maintain AIA cell identity and speculate that other transcriptional corepressors similarly function in the maintenance of specific cell identities and do so by silencing undesired gene expression through repression of transcriptional activators, such as EGL-13. Such a mechanism could explain how the breadth of transcriptional activation by terminal selectors can be fine-tuned in a coordinated fashion to fit the requirements of specific cell types, with selective transcriptional silencing providing a crucial aspect of proper cell-identity maintenance.

### CTBP-1 might physically interact with EGL-13 to maintain the AIA cell identity

The mammalian CtBPs (CtBP1 and CtBP2) bind PXDLS-like motifs on a number of diverse transcription factors to target specific genetic loci for silencing [49–51]. The mammalian ortholog of EGL-13, SOX6, interacts with the mammalian ortholog of CTBP-1, CtBP2, through a PLNLS motif located in SOX6 to repress *Fgf-3* expression in the developing mouse auditory otic vesicle [74]. This motif is 100% conserved in *C. elegans* EGL-13. We speculate that CTBP-1 interacts with EGL-13 through its PLNLS motif to regulate EGL-13 activity as part of AIA cell-identity maintenance. Specifically, we propose (Fig. 8) that CTBP-1 physically interacts with EGL-13 to target specific genetic loci for silencing as an aspect of normal AIA cell-identity maintenance. Following the establishment of the AIA cell fate, for which CTBP-1 is not required, CTBP-1 binds EGL-13, recruiting CTBP-1 to EGL-13 DNA binding sites. CTBP-1 then silences surrounding genetic loci, resulting in the repression of specific target genes (Fig. 8B). This repression is necessary for proper maintenance of the AIA cell identity, and when disrupted, as in *ctbp-1* mutants, CTBP-1 binding partners, such as EGL-13, inappropriately act as transcriptional activators in the AIAs, resulting in disruption of AIA gene expression, morphology and function (Fig. 8C). In the absence of such CTBP-1 interactors, as in *egl-13 ctbp-1* double mutants, aberrant transcription is not activated and some of the defects in AIA maintenance are avoided (Fig. 8D).

**Figure 8.**
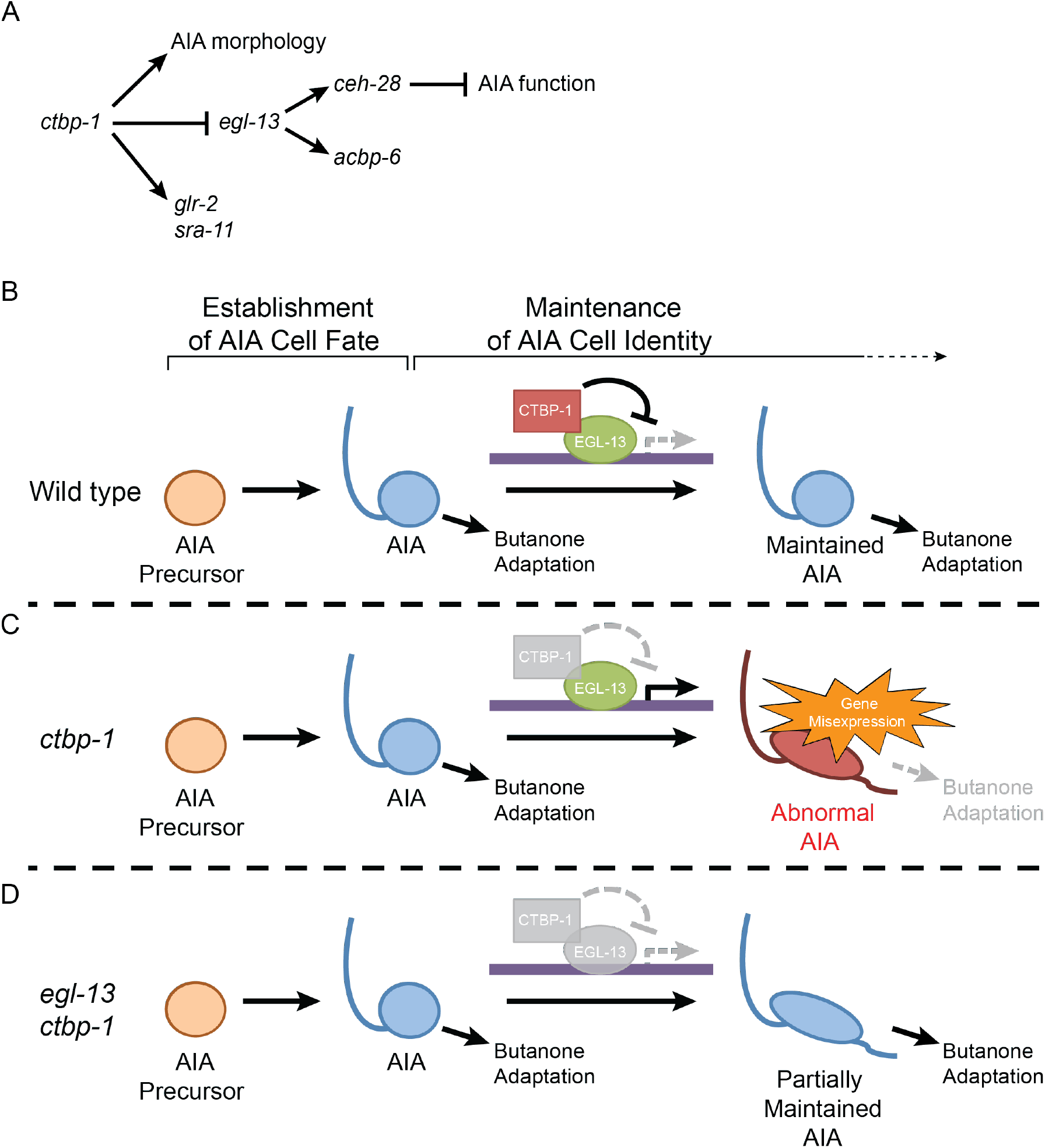
Model for the maintenance of the AIA cell identity by *ctbp-1*. (A) The genetic pathway in which *ctbp-1* promotes AIA morphology and *glr-2* and *sra-11* expression. *ctbp-1* also inhibits *egl-13*, thereby repressing expression of *ceh-28* and *acbp-6* in the AIAs and promoting proper AIA function (B-D) Model for how CTBP-1 maintains the AIA cell identity. (B) We propose that CTBP-1 acts in the maintenance but not establishment of the AIA cell identity, and does so by targeting specific genetic loci for regulation through physical interaction with transcription factors such as EGL-13. (C) In the absence of CTBP-1, EGL-13 and other CTBP-1 targets drive expression at multiple genetic loci, resulting in changes to the gene expression, morphology and function (as assessed by butanone adaptation) of the AIAs. (D) When EGL-13 activity is also removed, gene expression and cellular function are no longer perturbed, while normal morphology is not restored, resulting in a “Partially Maintained AIA.”

### CTBP-1 likely utilizes additional transcription factors besides EGL-13 to maintain the AIA cell identity

Our understanding of how CTBP-1 acts to maintain the AIA cell identity is incomplete. While we have identified a few genes with expression that changes in the absence of *ctbp-1* (*ceh-28*, *acbp-6*, *sra-11*, *glr-2*), none of these genes seems to individually account for the full range of AIA defects seen in older *ctbp-1* mutants. We speculate that there are many more unidentified transcriptional changes occurring in *ctbp-1* mutant AIAs that contribute to the observed AIA morphological and functional defects.

Our observations suggest that EGL-13 is not the sole transcription factor through which CTBP-1 functions to maintain AIA cell identity – neither AIA morphological defects nor some AIA gene-expression defects (i.e. *sra-11* and *glr-2* expression) in *ctbp-1* mutants were suppressed in *egl-13 ctbp-1* double mutants (Figs. 6D-G; 7A-B). We propose that CTBP-1 maintains different aspects of AIA cell identity through interactions with multiple different transcription factors. Given CTBP-1’s known function as a transcriptional corepressor and EGL-13’s observed role in driving gene misexpression in the absence of *ctbp-1*, we speculate that CTBP-1 likely utilizes not just EGL-13 but also other transcription factors (possibly through interacting with PXDLS-like motifs located in those transcription factors) to target multiple specific DNA sequences for transcriptional silencing, effectively turning these transcription factors into transcriptional repressors. When *ctbp-1* is absent, these unregulated transcription factors can aberrantly function as transcriptional activators, resulting in either the direct or indirect expression of genes which can in turn lead to defects in other aspects of cell identity. Such a mechanism for the selective and continuous silencing of multiple genetic loci in cell-type specific contexts by a transcriptional corepressor like CTBP-1 might explain how the broad activating activities of terminal selectors are restricted in the context of maintaining the identities of distinct cell types.

### CTBP-1 likely maintains the identities of other cells besides that of the AIAs

Others have previously reported a near pan-neuronal expression pattern of *ctbp-1* in *C. elegans* [53], suggesting that *ctbp-1* might be acting in more cells than just the AIAs to maintain cell identities. Why then have we thus far only been able to identify defects in the maintenance of the AIA identity in *ctbp-1* mutants? We speculate that, like the relatively subtle defects we have observed in AIA gene expression, morphology and function, *ctbp-1* mutant defects in the maintenance of other cell identities might be similarly subtle and easily missed if not specifically sought. In addition, the AIAs might be particularly susceptible to perturbations of maintenance of their identity, with defects manifesting either earlier in the life of the cell or in more distinct ways (e.g. more gene misexpression).

Both our findings and the work of others [54, 55] provide further support for the hypothesis that *ctbp-1* maintains other cell identities besides that of the AIAs. We observed that *ctbp-1* mutants have an additional AIA-independent chemotaxis defect (Figs. 4B-E; S5A-B), suggesting that other cells, likely neurons that sense and/or execute responses to volatile odors, are also dysfunctional. Additionally, others have shown that, in *ctbp-1* mutants another pair of *C. elegans* neurons, the SMDDs, display late-onset morphological abnormalities coupled with a defect in *C. elegans* foraging behavior associated with these cells [54, 55], indicating that CTBP-1 might act to maintain SMDD cell identity as well. The broad expression of *ctbp-1* throughout much of the *C. elegans* nervous system is also consistent with the hypothesis that *ctbp-1* functions broadly to maintain multiple neuronal cell identities [53].

### Transcriptional corepressors might function broadly in the maintenance of cell identities

The neuron-specific expression of *ctbp-1* [53] suggests that CTBP-1 likely does not function in maintaining the identities of non-neuronal cells. How might non-neuronal cell identities be maintained? We speculate that transcriptional corepressors function in maintaining cell identities in both neuronal and non-neuronal cells. There are known tissue-specific activities of other corepressor complexes, such as those of NCoR1 in mediating the downstream effects of hormone sensation in the mammalian liver [75, 76] or of <u>T</u>ransducin-<u>L</u>ike <u>E</u>nhancer of Split (TLE) in regulating gene expression and chromatin state in the developing mouse heart and kidney [77–79]. We propose that, by analogy to CTBP-1, distinct transcriptional corepressors might specialize in the maintenance of a wide range of cell identities in distinct tissue types throughout metazoa.

## Materials and Methods

### *C. elegans* strains and transgenes

All *C. elegans* strains were grown on Nematode Growth Medium (NGM) plates seeded with *E. coli* OP50 as described previously [80]. We used the N2 Bristol strain as wild type. Worms were grown at 20°C unless otherwise indicated. Standard molecular biology and microinjection methods, as previously described [81], were used to generate transgenic worms.

The following strains were used in this study:

| Strain | Genotype |
| --- | --- |
| N2 | wild type |
| MT15670 | *nls175*[*P<sub>ceh-28</sub>::gfp*] |
| MT15672 | *nls177*[*P<sub>ceh-28</sub>::gfp*] |
| MT15677 | *nls175; ctbp-1*(n4778) |
| MT16225 | *nls175; ctbp-1*(n4784) |
| MT15688 | *nls175; ctbp-1*(n4789) |
| MT15801 | *nls175; ctbp-1*(n4800) |
| MT15805 | *nls175; ctbp-1*(n4804) |
| MT15806 | *nls175; ctbp-1*(n4805) |
| MT15809 | *nls175; ctbp-1*(n4808) |
| MT15811 | *nls175; ctbp-1*(n4810) |
| MT15813 | *nls175; ctbp-1*(n4813) |
| MT15820 | *nls175; ctbp-1*(n4819) |
| MT15824 | *nls175; ctbp-1*(n4823) |
| MT15825 | *nls175; ctbp-1*(n4824) |
| MT15841 | *nls177; ctbp-1*(n4840) |
| MT15850 | *nls177; ctbp-1*(n4849) |
| MT15853 | *nls177; ctbp-1*(n4852) |
| MT15862 | *nls177; ctbp-1*(n4861) |
| MT15865 | *nls177; ctbp-1*(n4864) |
| MT15866 | *nls177; ctbp-1*(n4865) |
| MT26446 | *nls175; ctbp-1*(tm5512) |
| MT15918 | *nls175* introgressed into CB4856 “Hawaiian” background |
| MT16295 | *nls177* introgressed into CB4856 background |
| MT26522 | *nls175; ctbp-1*(n4784) introgressed into CB4856 background |
| MT23360 | *nls175; ctbp-1*(n4784); *nEx2346*[*ctbp-1*(+)] |
| MT23361 | *nls175; ctbp-1(n4784); nEx2347[ctbp-1(+)]* |
| MT23714 | *nls175; ctbp-1(n4784); nls743[P<sub>gcy-28.d</sub>::ctbp-1(+)]* |
| MT25271 | *nls843[P<sub>gcy-28.d</sub>::mCherry]* |
| MT26437 | *nls175; ctbp-1(n4784); nls843* |
| MT23365 | *nls175; ctbp-1(n4784); nEx2351[P<sub>hsp-16.2</sub>::ctbp-1(+); P<sub>hsp-16.41</sub>::ctbp-1(+)]* |
| MT18778 | *nls348[P<sub>ceh-28</sub>::mCherry]; lin-15AB(n765ts)* |
| MT20844 | *nls348[P<sub>ceh-28</sub>::mCherry]; ctbp-1(n4784)* |
| NH2466 | *ayls4[P<sub>egl-17</sub>::gfp]; dpy-20(e1282ts)* |
| MT26417 | *ayls4; nls348; ctbp-1(n4784)* |
| BW1946 | *ctls43[P<sub>dbl-1</sub>::gfp] unc-42(e270)* |
| MT23726 | *nls348; ctls43 unc-42(e270); ctbp-1(n4784)* |
| MT20852 | *nls491[P<sub>ser-7.b</sub>::mCherry]* |
| MT23427 | *nls491; ctbp-1(n4784)* |
| NY2080 | *ynls80[P<sub>flp-21</sub>::gfp]* |
| MT23718 | *nls348; ctbp-1(n4784); ynls80* |
| OH10237 | *otls326[P<sub>ins-1</sub>::gfp]* |
| MT26422 | *ctbp-1(n4784); otls326* |
| JN1716 | *pels1716[P<sub>ins-1s</sub>::gfp; P<sub>ttx-3</sub>::mCherry]* |
| MT23717 | *nls348; ctbp-1(n4784); pels1716* |
| OH11030 | *otls317[P<sub>mgl-1</sub>::mCherry]; otls379[P<sub>cho-1</sub>::gfp]* |
| MT26421 | *nls348; ctbp-1(n4784); otls317; otls379* |
| MT26420 | *ctbp-1(n4784); otls317* |
| MT25268 | *nls840[P<sub>gcy-28.d</sub>::gfp]* |
| MT25270 | *nls842[P<sub>gcy-28.d</sub>::gfp]* |
| MT26412 | *nls348; ctbp-1(n4784); nls840* |
| MT26438 | *nls348; ctbp-1(n4784); nls743; nls840* |
| MT26439 | *nls348; ctbp-1(n4784); nls840; nEx2351* |
| JN580 | *pels580[P<sub>ins-1s</sub>::casp1; P<sub>ins-1s</sub>::venus; P<sub>unc-122</sub>::gfp]* |
| MT23746 | *nls175; egl-13(n5937) ctbp-1(n4784)* |
| MT24129 | *nls175; egl-13(n6013) ctbp-1(n4784)* |
| MT25352 | *nls175; egl-13(n6313) ctbp-1(n4784)* |
| MT26486 | *nls175; egl-13(n5937) ctbp-1(n4784); nEx3062[egl-13(+)]* |
| MT26487 | *nls175; egl-13(n5937) ctbp-1(n4784); nEx3063[egl-13(+)]* |
| MT26549 | *nls175; egl-13(n6013) ctbp-1(n4784); nEx3080[egl-13(+)]* |
| MT26523 | *nls175; egl-13(n6313) ctbp-1(n4784); nEx3074[egl-13(+)]* |
| MT26548 | *nls175; egl-13(n6313) ctbp-1(n4784); nEx3079[egl-13(+)]* |
| MT26481 | *nls175; egl-13(n5937) ctbp-1(n4784); nEx3055[P<sub>gcy-28.d</sub>::egl-13(+)]* |
| MT26441 | *nls175; egl-13(n5937) ctbp-1(n4784); nls840* |
| MT26415 | *evls111[P<sub>rgef-1</sub>::gfp]; nls843* |
| MT26416 | *ctbp-1(n4784); evls111; nls843* |
| MT26444 | *otls123[P<sub>sra-11</sub>::gfp]; nls843* |
| MT26445 | *nls348; ctbp-1(n4784); otls123* |
| MT26524 | *nls348; egl-13(n5937) ctbp-1(n4784); otls123* |
| MT26504 | *nls843; ivEx138[P<sub>glr-2</sub>::gfp]* |
| MT26505 | *nls348; ctbp-1(n4784); ivEx138* |
| MT26550 | *nls348; egl-13(n5937) ctbp-1(n4784); ivEx138* |
| MT26581 | *nls843; nEx3081[P<sub>acbp-6</sub>::gfp]* |
| MT26551 | *nls348; ctbp-1(n4784); nEx3081* |
| MT26582 | *nls348; egl-13(n5937) ctbp-1(n4784); nEx3081* |
| MT26605 | *acbp-6(tm2995); nls175; ctbp-1(n4784)* |
| MT23725 | *nls175; ctbp-1(n4784) ceh-28(cu11)* |

### Plasmid construction

The *nIs175[P_ceh-28_::gfp]*, *nIs177[P_ceh-28_::gfp]* and *nIs348[P_ceh-28_::mCherry]* transgenes have been previously described [59]. *nIs743[P_gcy-28.d_::ctbp-1(+)]* contains 3.0 kb of the 5’ promoter of *gcy-28.d* fused to the *ctbp-1a* coding region inserted into plasmid pPD49.26. *nIs840[P_gcy-28.d_::gfp]* contains 3.0 kb of the 5’ promoter of *gcy-28.d* inserted into pPD95.77. *nIs843[P_gcy-28.d_::mCherry]* contains 3.0 kb of the 5’ promoter of *gcy-28.d* inserted into pPD122.56 containing mCherry. *nEx2351[P_hsp-16.2_::ctbp-1(+);P_hsp-16.41_::ctbp-1(+)]* contains *ctbp-1a* cDNA, isolated by RT-PCR, inserted into pPD49.73 and pPD49.83. *nEx3055[P_gcy-28.d_::egl-13(+)]* contains 3.0 kb of the 5’ promoter of *gcy-28.d* fused to the *egl-13* coding region inserted into pPD49.26. *nEx3081[P_acbp-6_::gfp]* contains 2.0 kb of the 5’ promoter of *acbp-6* inserted into pPD122.56. Plasmid construction was performed using Infusion cloning enzymes (Takara Bio, Mountain View, CA).

### Mutagenesis screens

*ctbp-1* mutants were isolated from genetic screens for mutations that cause the survival of the M4 sister cell as scored by extra GFP-positive cells carrying the M4-cell-specific markers *nIs175[P_ceh-28_::gfp]* or *nIs177[P_ceh-28_::gfp]* [58, 59]. *egl-13* mutants were isolated from genetic screens for mutations that suppress *nIs175* misexpression in the AIAs of *ctbp-1(n4784)* mutants while retaining GFP expression in the M4 neuron. For both screens, mutagenesis was performed with ethyl methanesulfonate (EMS) as previously described [80]. Mutagenized P_0_ animals were allowed to propagate, and their F_2_ progeny were synchronized by hypochlorite treatment and screened at the L4 stage for extra GFP-positive cells (*ctbp-1* screens) or fewer GFP-positive cells (suppressor screens) on a dissecting microscope equipped to examine fluorescence. From both screens, mutant alleles were grouped into functional groups by complementation testing when possible. Mutants were mapped using SNP mapping [70] by crossing mutants to strains containing *nIs175*, *nIs177*, or *nIs175;ctbp-1(n4784)* introgressed into the Hawaiian strain CB4856. Whole-genome sequencing was performed on mutants and a combination of functional groupings and mapping data suggested genes with mutations that were likely causal for the mutant phenotypes. Rescue of mutant phenotypes with wild-type *ctbp-1(+)* and *egl-13(+)* constructs as well as the mutant phenotype of a separately isolated deletion allele of *ctbp-1, tm5512,* confirmed the identities of the causal mutations.

### Microscopy

All images were obtained using an LSM 800 confocal microscope (Zeiss) and ZEN software. Images were processed and prepared for publication using FIJI software and Adobe Illustrator.

### Heat-shock assays

Rescue of AIA defects in day 1 adult worms was assayed using the *nEx2351[P_hsp-16.2_::ctbp-1;P_hsp-16.41_::ctbp-1]* transgene. Worms were synchronized and grown at 20°C. Subsets of L1 and L4 worms carrying *nEx2351* were removed from this population for scoring at the appropriate stages. At the L4 stage, half of the worms were heat-shocked at 34°C for 30 minutes and returned to 20°C for 24 hours while the other half remained at 20°C throughout. After 24 hours, heat-shocked and non-heat-shocked worms carrying *nEx2351* were scored.

### Single-cell RNA-sequencing

#### Dissociation of animals into cell suspensions

Single-cell suspensions were generated as described [82–84] with minor modifications. Briefly, synchronized populations of worms were grown on NGM plates seeded with OP50 to the L4 larval stage. Worms were harvested from these plates, washed three times with M9 buffer and treated with SDS-DTT (200 mM DTT, 0.25% SDS, 20 mM HEPES, 3% sucrose, pH 8.0) for two to three minutes. Worms were washed five times with 1x PBS and treated with pronase (15 mg/mL) for 20-23 minutes. During the pronase treatment, worm suspensions were pipetted with a P200 pipette rapidly for four sets of 80 repetitions. The pronase treatment was stopped by the addition of L-15-10 media (90% L-15 media, 10% FBS). The suspension was then passed through a 35 µm nylon filter into a collection tube, washed once with 1x PBS, and prepared for FACS.

#### FACS of fluorescently-labeled neurons

FACS was performed using a BD FACSAria III cell sorter running BD FACS Diva software. DAPI was added to samples at a final concentration of 1 µg/mL to label dead and dying cells. GFP-positive, DAPI-negative neurons were sorted from the single-cell suspension into 1x PBS containing 1% FBS. Non-fluorescent and single-color controls were used to set gating parameters. Cells were then concentrated and processed for single-cell sequencing.

#### Single-cell sequencing

Samples were processed for single-cell sequencing using the 10X Genomics Chromium 3’mRNA-sequencing platform. Libraries were prepared using the Chromium Next GEM Single Cell 3’ Kit v3.1 according to the manufacturer’s protocol. The libraries were sequenced using an Illumina NextSeq 500 with 75 bp paired end reads.

#### Single-cell RNA-sequencing data processing

Data processing was performed using 10X Genomics’ CellRanger software (v4.0.0). Reads were mapped to the *elegans* reference genome from Wormbase, version WBcel235. For visualization and analysis of data, we used 10X Genomics’ Loupe Browser (v4.2.0). AIAs were identified by expression of multiple AIA markers confirmed to be expressed in both wild-type and *ctbp-1* mutant AIAs (i.e. *gcy-28, ins-1, cho-1*; Fig. 2B). Candidate genes for misexpression (either ectopic or missing) in mutant AIAs were identified and tested as described in the text.

### Morphology scoring

We assayed AIA morphology by visualizing and imaging AIAs expressing *nIs840* using an LSM 800 confocal microscope (Zeiss) and a 63x objective. AIA cell body length and area were quantified using FIJI software.

### Image blinding and scoring

A subset of 60 wild-type and 60 *ctbp-1* mutant images per stage (randomly chosen from the existing images taken to measure AIA cell body length) were selected and the genotype of each was blinded. Blinded images were then scored as either “Normal” or “Elongated” in appearance in batches of 40 images (20 each of wild-type and *ctbp-1* mutant, randomly assorted), repeated three times per stage. Scored images were then matched back to their genotypes and percentage of AIAs scored as “Elongated” per genotype was calculated and graphed.

### Behavioral Assays

#### Butanone adaptation

Assay conditions were adapted from Cho et al., 2016 [66]. Staged worms were washed off non-crowded NGM plates seeded with *E. coli* OP50 with S basal. Worms were washed two times with S basal and split evenly into the “naïve” and “conditioned” populations. Naïve worms were incubated in 1 mL S basal for 90 minutes. Conditioned worms were incubated in 1 mL S basal with 2-butanone diluted to a final concentration of 120 µM for 90 minutes. During conditioning, unseeded NGM plates were spotted with two 1 µL drops of 10% ethanol (“control”) and two 1 µL drops of 2-butanone diluted in 10% ethanol at 1:1000 (“odor”) as well as four 1 µL drops of 1 M NaN_3_ at the same loci. After conditioning, both populations were washed three more times in S basal and placed at the center of the unseeded NGM plates. Worms were allowed to chemotax for two hours. Plates were moved to 4°C for 30-60 minutes to stop the assay and then scored. Worms that had left the origin were scored as chemotaxing to the odor spots (“#odor”) or control spots (“#control”), and a chemotaxis index was determined as (#odor - #control) / (#odor + #control). Assays were repeated on at least three separate days with one to three plates per strain ran in parallel on any given day based on the number of appropriately-staged worms available. Plates in which fewer than 50 worms left the origin were not scored.

#### Chemotaxis assays

L4 worms were washed off non-crowded NGM plates seeded with *E. coli* OP50 with S basal. Worms were washed three times with S basal. Unseeded NGM plates were spotted with two 1 µL drops of 100% ethanol (“control”) and two 1 µL drops of diacetyl diluted in 100% ethanol at 1:1000 or two 1 µL drops of isoamyl alcohol diluted in 100% ethanol at 1:100 (“odor”) as well as four 1 µL drops of 1 M NaN_3_ at the same loci. Worms were placed at the center of the unseeded NGM plates. Worms were allowed to chemotax for two hours. Plates were moved to 4°C for 30-60 minutes to stop the assay and then scored. Worms that had left the origin were scored, and a chemotaxis index was determined as above. Assays were repeated on at least three separate days. Plates in which fewer than 40 worms left the origin were not scored.

### Statistical analyses

Unpaired t-tests were used for the comparisons of AIA gene expression and AIA morphological features. One-way ANOVA tests with Tukey’s correction were used for comparisons of AIA morphological features and for behavioral assays. Statistical tests were performed using GraphPad Prism software (GraphPad Prism version 6.0h, RRID: SCR_002798).

### Accession Number

The GEO accession number for the RNA-Seq dataset in this paper is GSE179484.

## Acknowledgments

We thank N. An, R. Droste, S. Mitani, and the *Caenorhabditis* Genetics Center (CGC), which is funded by NIH Office of Research Infrastructure Programs (P40 OD010440), for strains and reagents. We thank C. Diehl, E. Lee, C. Pender, S. Sando, V. Dwivedi, K. Burkhart, S. Wong, C. Fincher, C. Cho, G. Johnson, D. Lee, D. Ghosh, A. Amon, P. Reddien, and Horvitz laboratory members for discussion and advice.

## Funding

**National Institutes of Health (GM024663)**

- Josh Saul
- H. Robert Horvitz

**National Institutes of Health (T32GM007287)**

- Josh Saul

**Howard Hughes Medical Institute**

- Josh Saul
- Takashi Hirose
- H. Robert Horvitz

**Friends of the McGovern Institute Fellowship (2733360)**

- Josh Saul

## Author Contributions

H.R.H. supervised the project. T.H. initiated the project. J.S. and T.H. designed and performed experiments, generated reagents, and analyzed data. All authors contributed to interpretation of data. J.S. wrote the original manuscript draft. All authors contributed to review and editing of the manuscript.

---

**Figure S1.**
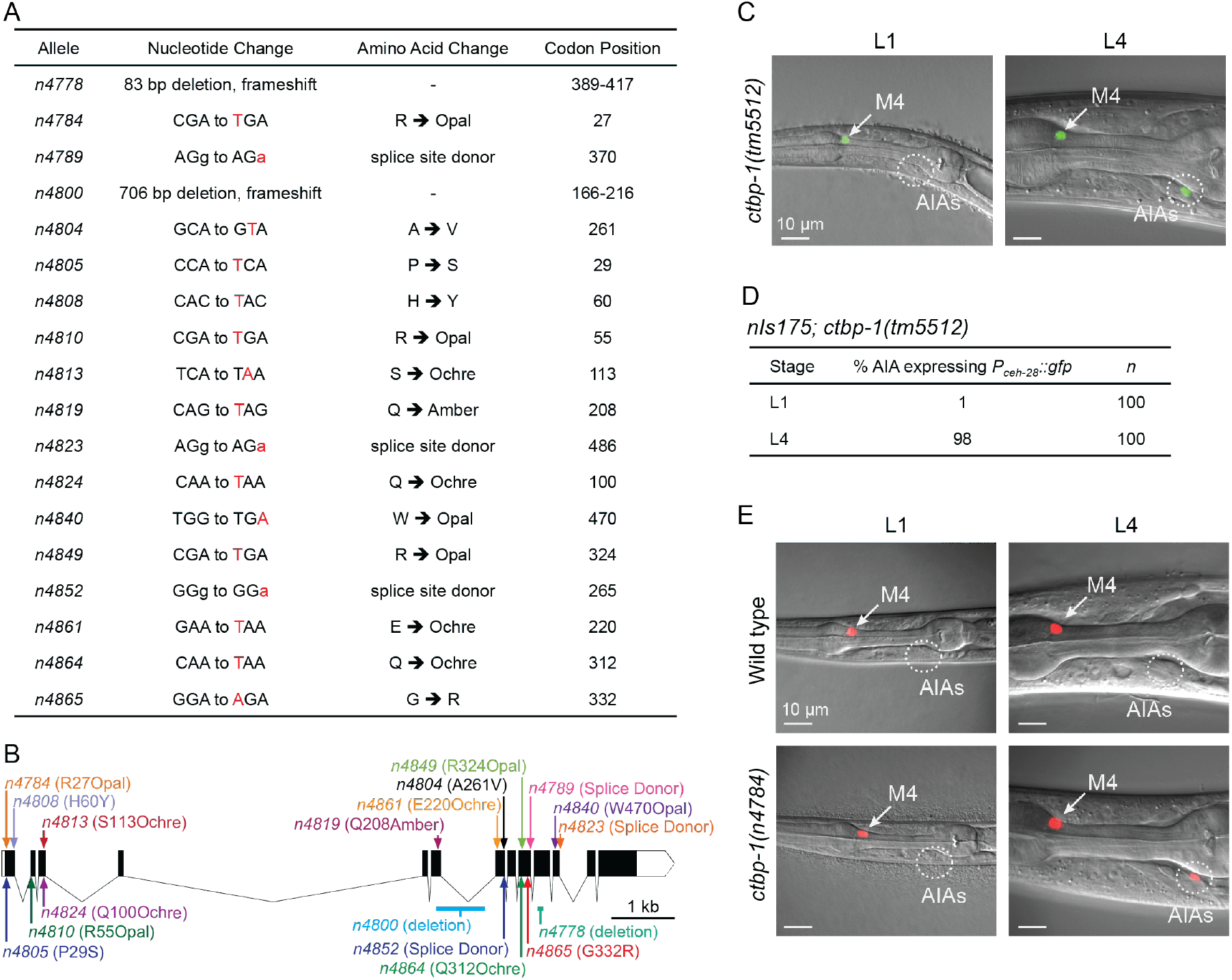
Additional *ctbp-1* mutant alleles cause misexpression of *P_ceh-28_::gfp* in the AIA neurons. (A) Table of *ctbp-1* mutant alleles we showed to result in *nIs175[P_ceh-28_::gfp]* or *nIs177[P_ceh-28_::gfp]* misexpression in the AIA neurons. Specific nucleotide changes are denoted in red. Codon positions correspond to the *ctbp-1a* isoform. (B) Gene diagram of the *ctbp-1a* isoform showing all 18 *ctbp-1* alleles isolated in this study. Arrows, point mutations. Lines, deletions. Scale bar (bottom right), 1 kb. (C) Expression of *nIs175* in *ctbp-1(tm5512)* L1 and L4 mutant worms. Arrow, M4 neuron. Circle, AIAs. Scale bar, 10 µm. (D) Percentage of *ctbp-1(tm5512)* worms expressing *nIs175* in the AIA neurons at the L1 and L4 larval stages. (E) *nIs348[P_ceh-28_::mCherry]* expression in wild-type (top) and *ctbp-1(n4784)* (bottom) worms at the L1 larval stage (left) and L4 larval stage (right). Arrow, M4 neuron. Circle, AIAs. Scale bar, 10 µm. All strains in Fig. S1A-C contain the either *nIs175[P_ceh-28_::gfp]* or *nIs177[P_ceh-28_::gfp]*. Images are oriented such that left corresponds to anterior, top to dorsal.

**Figure S2.**
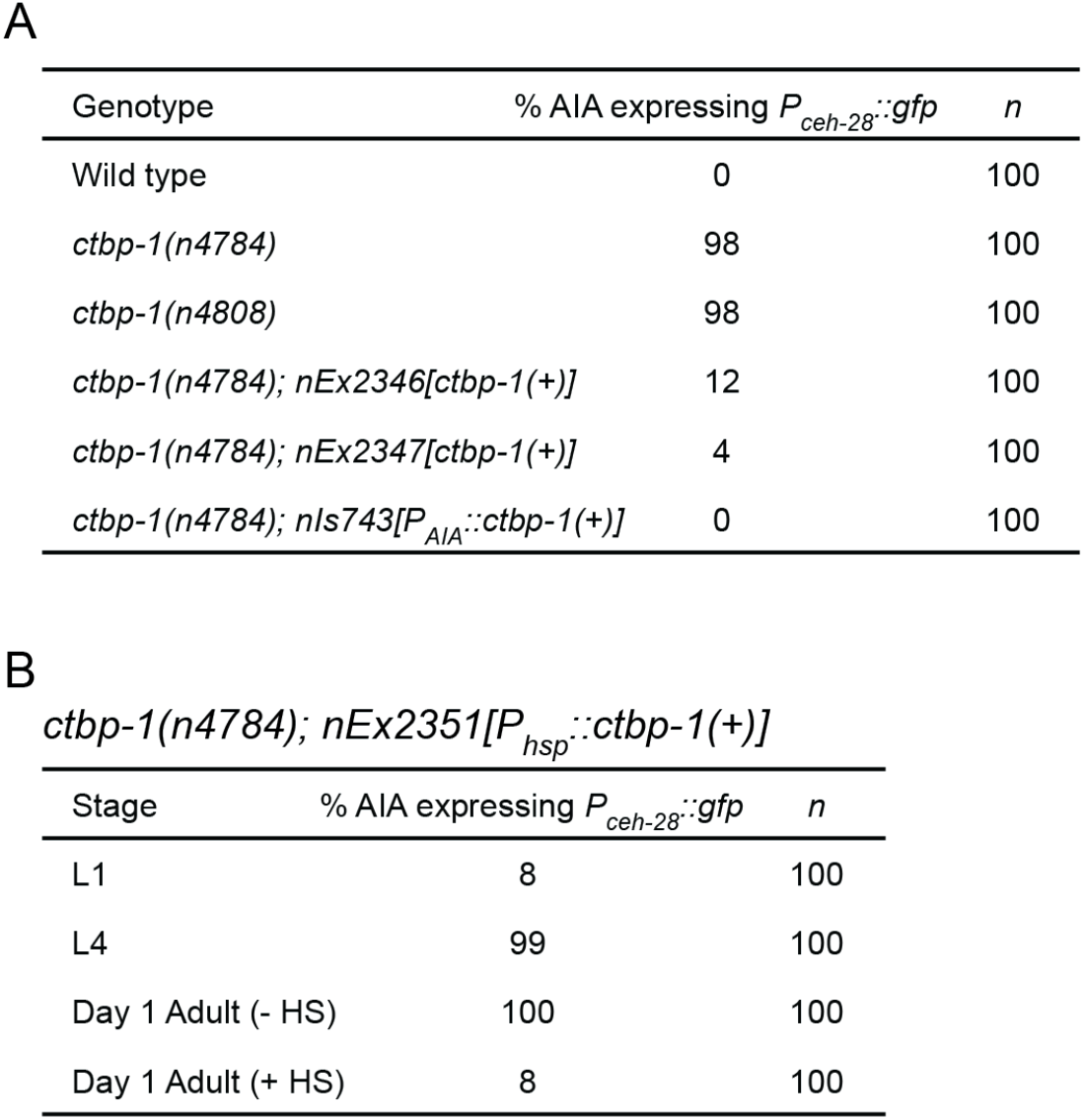
Quantification of *ctbp-1* strains misexpressing *P_ceh-28_::gfp*. (A) Percentage of worms of indicated genotypes expressing *nIs175[P_ceh-28_::gfp]* in the AIA neurons at the L4 larval stage. (B) Percentage of *ctbp-1(n4784); nEx2351[P_hsp-16.2_::ctbp-1(+);P_hsp-16.41_::ctbp-1(+)]* worms expressing *nIs175* at the L1, L4 and day 1 adult stages. Day 1 adults shown ± heat shock (HS) at the L4 stage.

**Figure S3.**
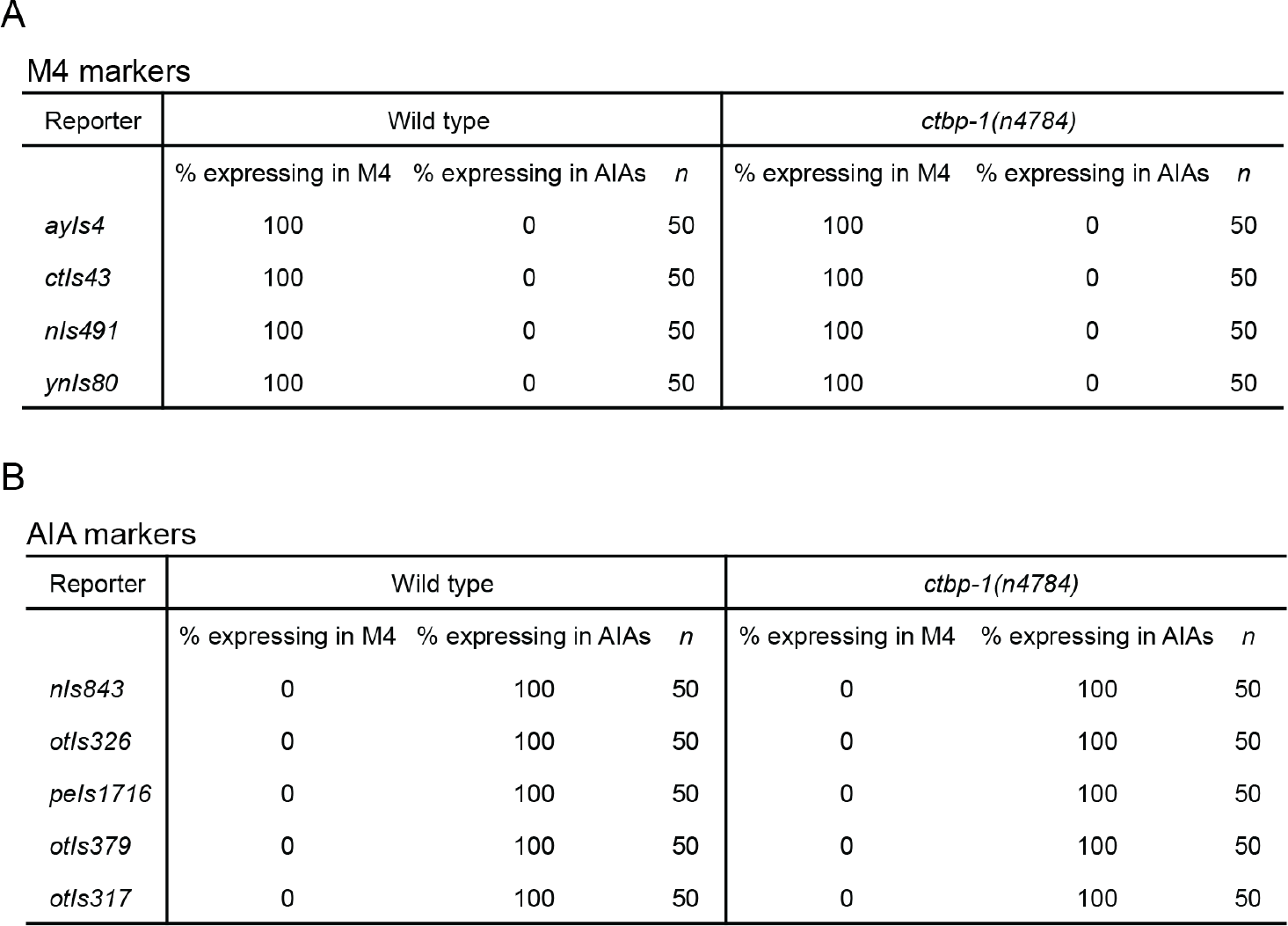
Quantification of M4 and AIA marker expression. (A-B) Quantification of wild-type and *ctbp-1(n4784)* L4 worms expressing the indicated (A) M4 or (B) AIA markers from Fig. 2A-B in the M4 and AIA neurons.

**Figure S4.**
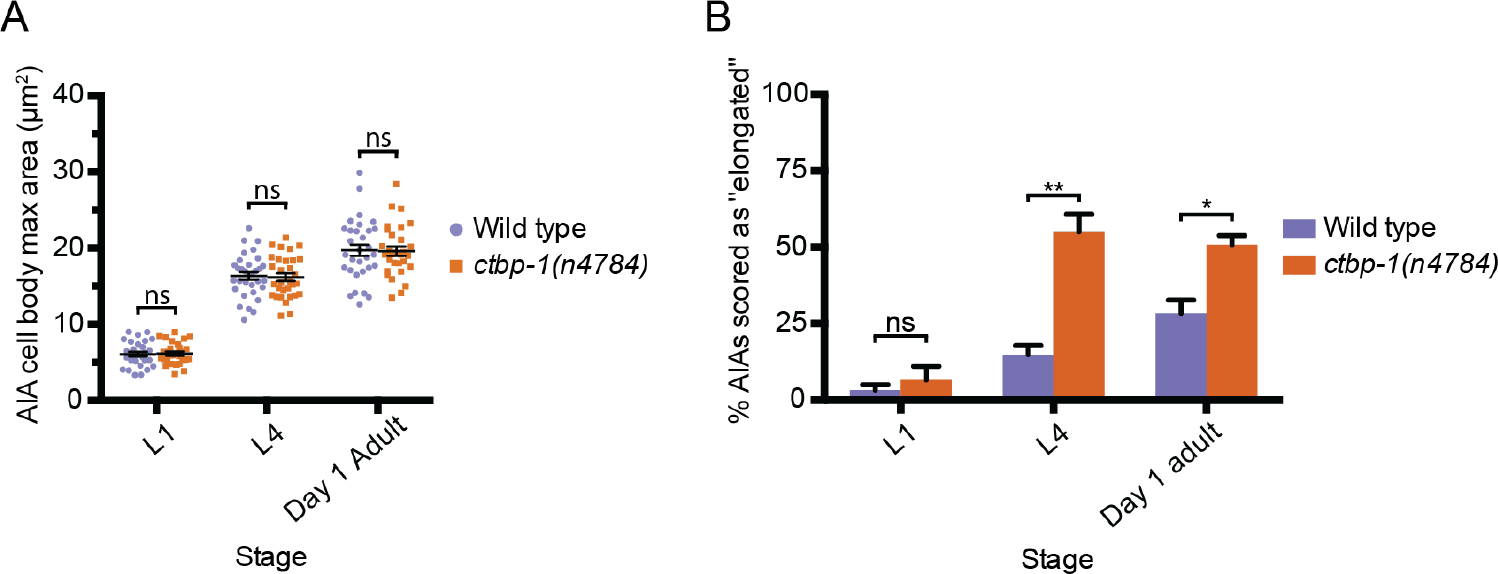
Loss of *ctbp-1* results in a disruption of AIA morphology but not AIA size. (A) Quantification of maximum AIA cell body area in wild-type and *ctbp-1(n4784)* worms at L1, L4 and day 1 adult stages. Both strains contain *nIs840[P_gcy-28.d_::gfp]* and the *ctbp-1* strain contains *nIs348[P_ceh-28_::mCherry]*. Mean ± SEM. *n* ≥ 30 AIAs scored per strain per stage. ns, not significant, unpaired t-test. (B) Scoring of wild-type and *ctbp-1(n4784)* AIA images at the L1, L4 and day 1 adult stages. A random subset of AIA images used for length measurements in Fig. 3D, 3H and 3L were blinded and scored as having either “Normal” or “Elongated” AIA cell bodies. *n* ≥ 20 AIAs scored per strain per stage, 3 replicates. ns, not significant, *p<0.05, **p<0.01, unpaired t-test.

**Figure S5.**
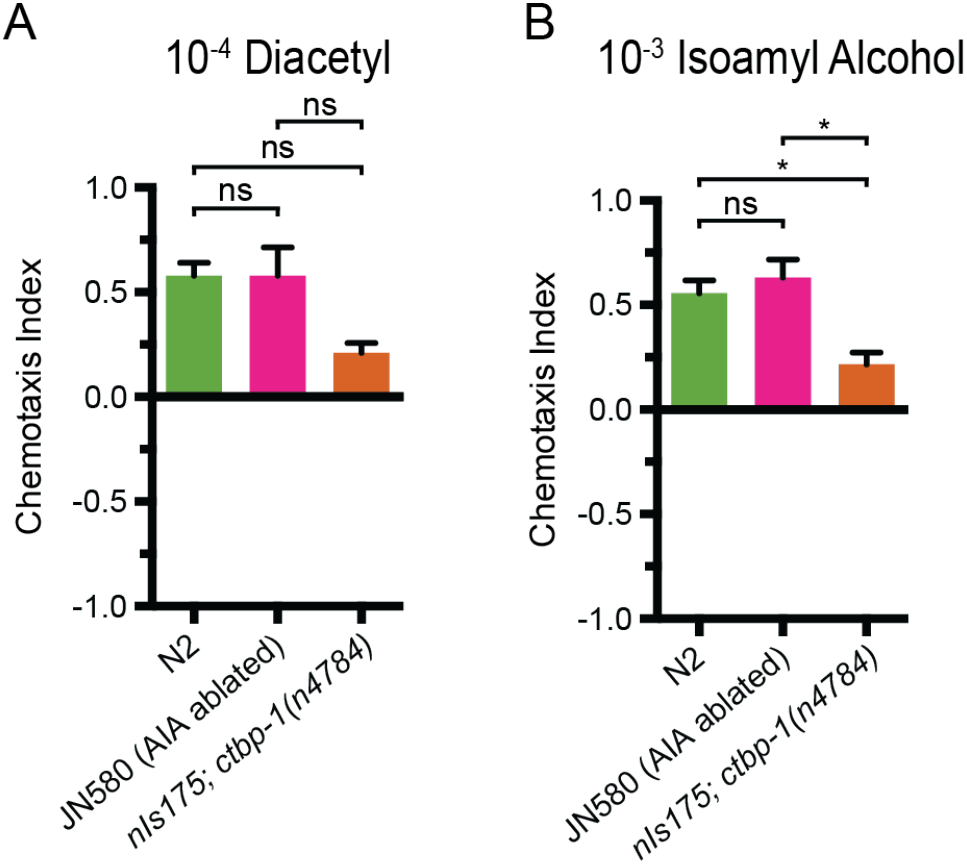
*ctbp-1* mutants display a non-AIA-dependent chemotaxis defect. (A-B) Chemotaxis indices of wild-type (N2), AIA-ablated (JN580) and *nIs175*; *ctbp-1(n4784)* mutants at the L4 larval stage to (A) diacetyl or (B) isoamyl alcohol diluted in pure ethanol. Mean ± SEM. *n* ≥ 3 assays per condition, ≥ 40 worms per assay. ns, not significant, *p<0.05, one-way ANOVA with Tukey’s correction.

**Figure S6.**
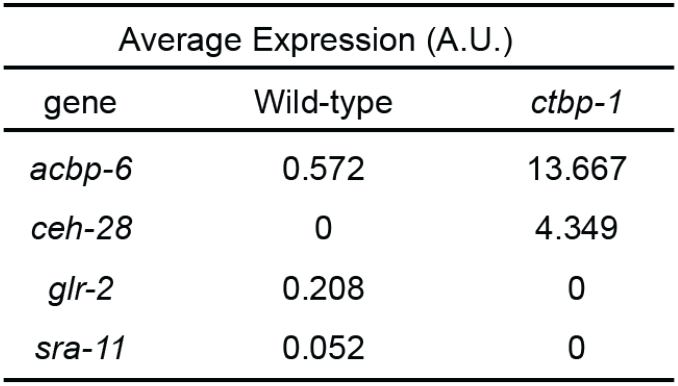
Gene expression in wild-type and *ctbp-1* AIAs. Average expression level of confirmed scRNA-Seq hits in the AIAs of L4 wild-type and *ctbp-1* mutant animals. A.U., arbitrary expression units.

**Figure S7.**
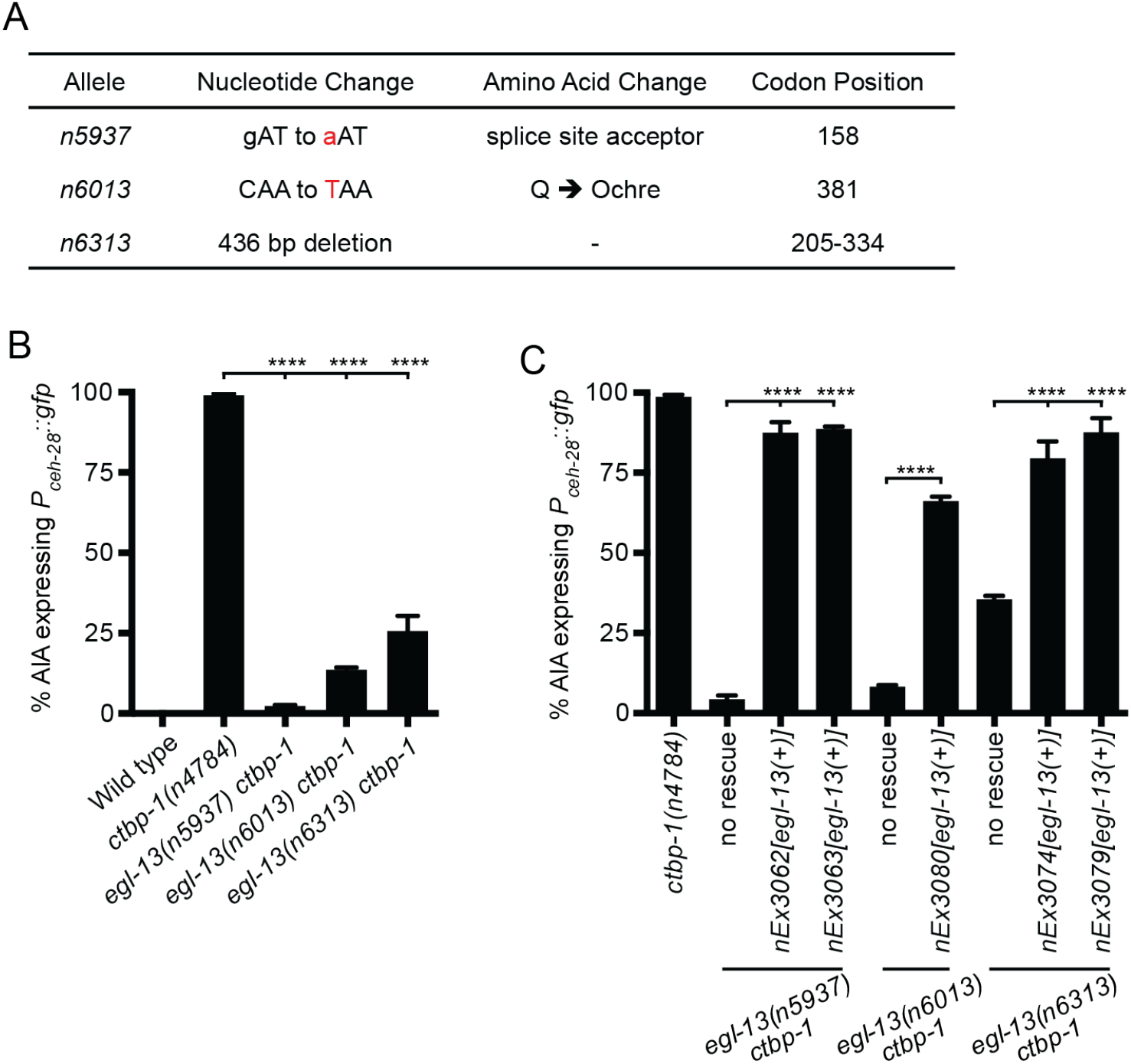
Characterization of *egl-13* alleles isolated as *ctbp-1* suppressors. (A) Table of *egl-13* mutant alleles isolated in this study as suppressors of *ctbp-1*-mediated *nIs175[P_ceh-28_::gfp]* misexpression in the AIA neurons. Specific nucleotide changes are denoted in red. Codon positions correspond to *egl-13a* isoform. (B) Percentage of wild-type, *ctbp-1(n4784)* and *egl-13 ctbp-1* worms expressing *nIs175* in the AIA neurons at the L4 larval stage. Mean ± SEM. *n* ≥ 100 worms scored per strain, 3 biological replicates. ****p<0.0001, one-way ANOVA with Tukey’s correction. (C) Percentage of *ctbp-1(n4784), egl-13 ctbp-1* and *egl-13 ctbp-1* worms carrying transgenic constructs expressing wild-type *egl-13* under its native promoter expressing *nIs175* in the AIA neurons at the L4 larval stage. Mean ± SEM. *n* ≥ 50 worms scored per strain, 3 biological replicates. ****p<0.0001, one-way ANOVA with Tukey’s correction. All strains in Fig. S7B-C contain *nIs175[P_ceh-28_::gfp]*.

**Figure S8.**
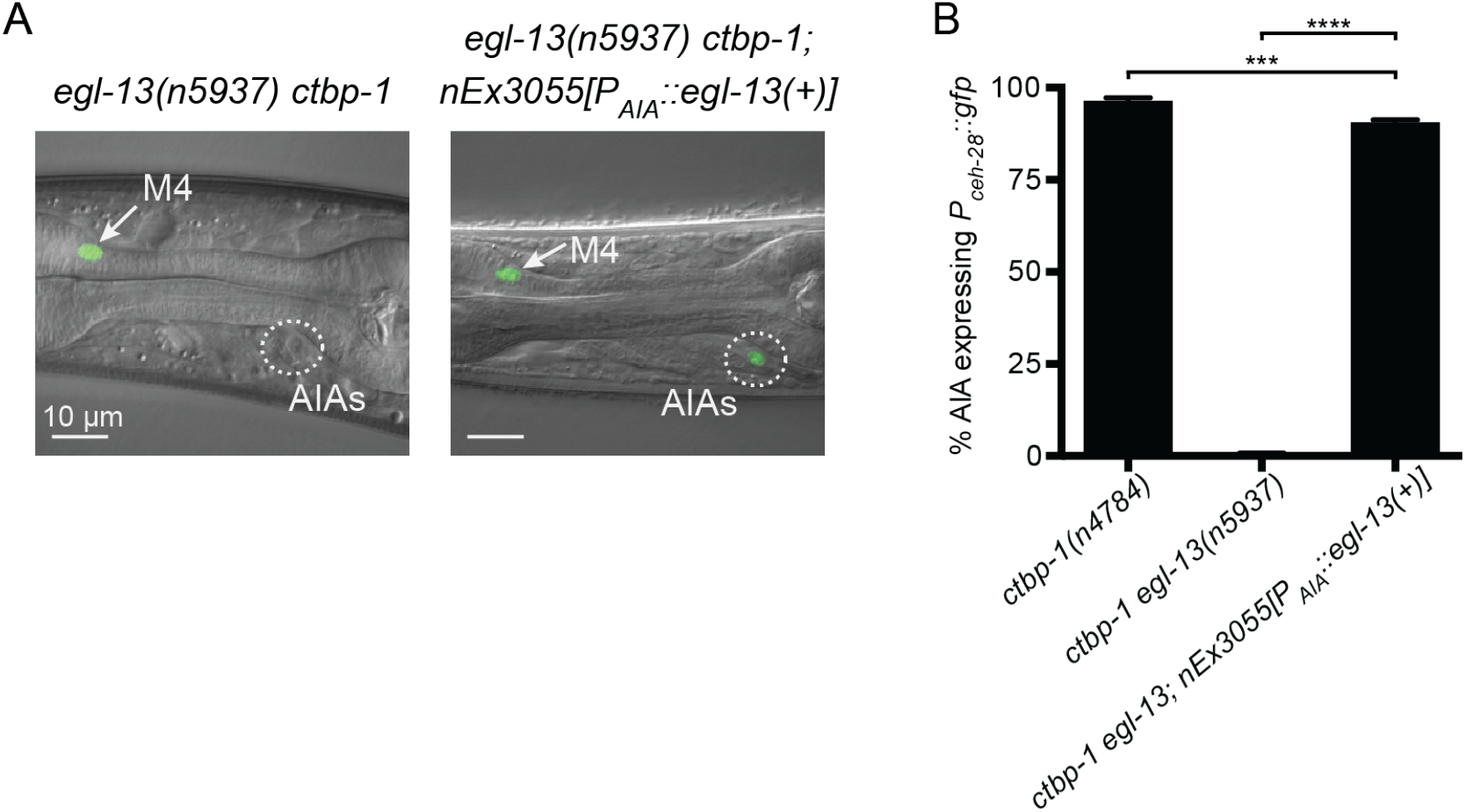
EGL-13 functions cell-autonomously to regulate AIA gene expression. (A) Expression of *nIs175[P_ceh-28_::gfp]* in *egl-13(n5937) ctbp-1(n4784)* (left panel), and *egl-13 ctbp-1* mutants carrying an extrachromosomal array expressing wild-type *egl-13* under the AIA-specific promoter *gcy-28.d* (*nEx3055*, right panel) in L4 worms. Arrow, M4 neuron. Circle, AIAs. Scale bar, 10 µm. (B) Percentage of *ctbp-1(n4784)*, *egl-13(n5937) ctbp-1* and *egl-13 ctbp-1; nEx3055* worms expressing *nIs175* in the AIA neurons at the L4 larval stage. All strains contain *nIs175[P_ceh-28_::gfp].* Mean ± SEM. *n* = 100 worms scored per strain, 3 biological replicates. ***p<0.001, ****p<0.0001, one-way ANOVA with Tukey’s correction. The alleles used for all panels of this figure were *ctbp-1(n4784*) and *egl-13(n5937)*. All strains contain *nIs175[P_ceh-28_::gfp]*. Images are oriented such that left corresponds to anterior, top to dorsal.

